# EM-seq: Detection of DNA Methylation at Single Base Resolution from Picograms of DNA

**DOI:** 10.1101/2019.12.20.884692

**Authors:** Romualdas Vaisvila, V. K. Chaithanya Ponnaluri, Zhiyi Sun, Bradley W. Langhorst, Lana Saleh, Shengxi Guan, Nan Dai, Matthew A. Campbell, Brittany S. Sexton, Katherine Marks, Mala Samaranayake, James C. Samuelson, Heidi E. Church, Esta Tamanaha, Ivan R. Corrêa, Sriharsa Pradhan, Eileen T. Dimalanta, Thomas C. Evans, Louise Williams, Theodore B. Davis

## Abstract

Bisulfite sequencing is widely used to detect 5mC and 5hmC at single base resolution. However, bisulfite treatment damages DNA resulting in fragmentation, loss of DNA and biased sequencing data. To overcome this, we developed Enzymatic Methyl-seq (EM-seq), an enzymatic based approach that uses as little as 100 pg of DNA. EM-seq outperformed bisulfite converted libraries in all metrics examined including coverage, duplication, sensitivity and nucleotide composition. EM-seq libraries displayed even GC distribution, improved correlation across input amounts as well as increased representation of genomic features. These data indicate that EM-seq is more accurate and reliable than whole genome bisulfite sequencing (WGBS).

## Background

There are approximately 2.6 billion cytosines in the human genome, and when both DNA strands are considered, 56 million of those are followed by guanines (CpGs) (1). In mammalian genomes, 70% to 80% of CpG are modified (2). Cytosines modified at the 5^th^ carbon position with a methyl group result in 5-methylcytosine (5mC) and oxidation of 5mC results in the formation of 5-hydroxymethylcytosine (5hmC). These modifications are important due to their impact on a wide range of biological processes including gene expression and development (3). Cytosine modifications are often linked with altered gene expression, for example, methylated cytosines are often associated with transcriptional silencing and are found at transcription start sites of repressed genes (4) or at repetitive DNA and transposons (5). Recently however, it has been reported that some genes can be activated by 3’ CpG island methylation during development (6). The ability to accurately detect 5mC and 5hmC can have profound implications in understanding biological processes and in the diagnosis of diseases such as cancer.

To date, bisulfite sequencing has been the accepted standard for mapping the methylome. Sodium bisulfite chemically modifies DNA and results in the conversion of unmethylated cytosines to uracils (Supplemental Figure 1), however, 5mC and 5hmC are not converted under these reaction conditions (7). Sequencing distinguishes cytosines from these modified forms as they are read as thymines and cytosines respectively (8). Despite its widespread use amongst epigenetic researchers, bisulfite sequencing also has significant drawbacks. It requires extreme temperatures and pH which causes depyrimidination of DNA resulting in DNA degradation (9). Furthermore, cytosines are damaged disproportionately compared to 5mC or 5hmC. As a result, sequencing libraries made from converted DNA have an unbalanced nucleotide composition. All of these issues taken together result in libraries with reduced mapping rates and skewed GC content representation, with a general underrepresentation of G- and C-containing dinucleotides and over-representation of AA-, AT- and TA-containing dinucleotides, when compared to a non-converted genome (10). Therefore, the damaged libraries do not adequately cover the genome, and can include many gaps with little or no coverage. Increasing the sequencing depth of these libraries may recover some missing information, but at steep sequencing costs.

These bisulfite library limitations have driven the development of new approaches for mapping 5mC and 5hmC, in combination or independently, for epigenome analysis. The methylation dependent restriction enzymes (MDRE), MspJI and AbaSI were used to detect 5mC or 5hmC throughout the genome (11, 12) and the use of AbaSI was further adapted for use in single cell 5hmC detection (13). These methods also have drawbacks that are related to the enzymatic properties of MDRE, such as variable target site cleavage efficiency that results in potential biases within data sets. Another enzymatic method, ACE-seq, uses two enzymes, T4-phage β glucosyltransferase (T4-βGT) and apolipoprotein B mRNA editing enzyme, catalytic polypeptide-like 3A (APOBEC3A). T4-βGT protects 5hmC against enzymatic deamination by APOBEC3A. Cytosines and 5mCs are deaminated to uracil and thymine respectively and then sequenced as thymines and 5hmCs are sequenced as cytosines. Comparing the reads to an unconverted genome enables the identification of 5hmCs (14). Recently, TET-assisted pyridine borane sequencing (TAPS), which combines an enzymatic and chemical reaction to detect 5mC and 5hmC was published (15).TET1 is used to oxidize 5mC and 5hmC to 5-carboxycytosine (5caC). The 5caCs are then reduced to dihydrouracil (DHU) using pyridine borane. Subsequent PCR converts DHU to thymine, enabling cytosines and the corresponding cytosine modifications to be differentiated. This method is potentially hindered by the difficulty to prepare hyperactive TET1, which is required for complete oxidization of 5mC to 5caC.

At this time, only destructive conversion methods are available to detect methylated cytosines. Here we describe Enzymatic Methyl-seq (EM-seq), the first method to use only enzymatic reactions for single base resolution mapping of 5mC/5hmC. Enzymatic Methyl-seq provides a much-needed alternative to bisulfite sequencing. This method utilizes two enzymatic steps to differentiate between cytosine and its modified forms, 5mC and 5hmC. The results described herein show that enzymatically converted DNA is not fragmented in the same way as bisulfite converted DNA resulting in less biased sequencing data, even genome coverage and increased detection of CpGs. The ability to obtain a complete methylome has been incredibly difficult to achieve using bisulfite sequencing. EM-seq opens up new avenues to investigate methylomes and gain a deeper understanding of development and disease.

## RESULTS

### Enzymatic Detection of 5mC and 5hmC on Genomic DNA

Enzymatic detection of 5mC and 5hmCs requires two sets of reactions. The first reaction uses ten-eleven translocation dioxygenase 2 (TET2) and T4 phage β-glucosyltranferase (T4-βGT). TET2 is a Fe(II)/alpha-ketoglutarate-dependent dioxygenase that catalyzes the oxidization of 5-methylcytosine to 5-hydroxymethylcytosine (5hmC), 5-formylcytosine (5fC), and 5-carboxycytosine (5caC) in three consecutive steps with the concomitant formation of CO_2_ and succinate (Figure 1A). T4-βGT catalyzes the glucosylation of the formed 5hmC as well as pre-existing genomic 5hmCs to 5-(β-glucosyloxymethyl)cytosine (5gmC) (Figure 1B). These reactions protect 5mC and 5hmC against deamination by APOBEC3A (Figure 1C, Figure 3C). This ensures that only cytosines are deaminated to uracils, thus enabling the discrimination of cytosine from its methylated and hydroxymethylated forms. The following sections characterize the catalytic actions of TET2, T4-βGT, and APOBEC3A.

**Figure 1.**
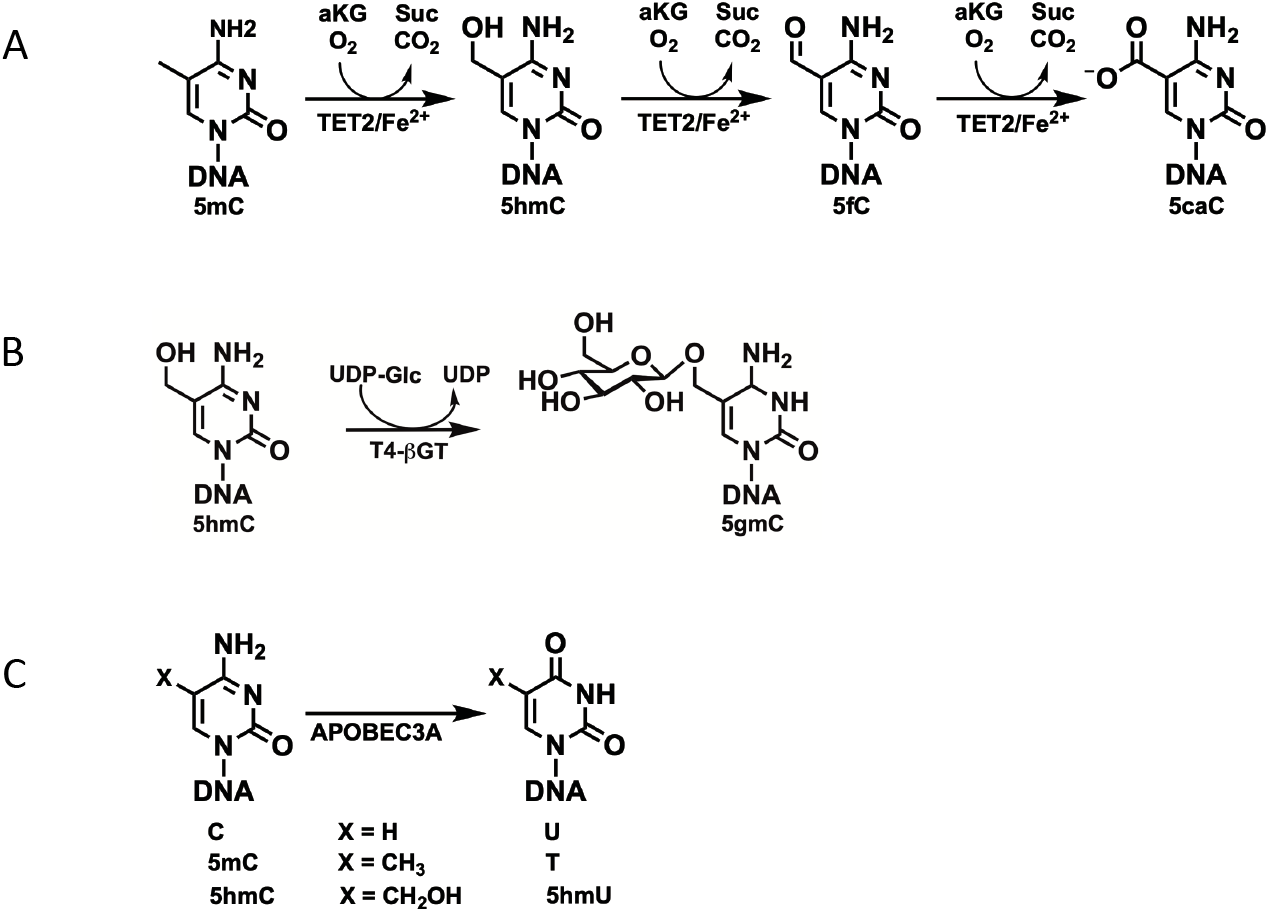
Enzymatic Activity of Enzymes Involved in Detection of 5mC and 5hmC. (A) TET2 catalyzes the oxidization of 5mC to 5hmC, 5fC, and 5caC in three consecutive steps. (B) The T4-phage enzyme T4-βGT glucosylates pre-existing genomic 5hmC as well as 5hmC formed by the action of TET2 to 5-(β-glucosyloxymethyl)cytosine (5gmC). (C) APOBEC3A (apolipoprotein B mRNA editing enzyme, catalytic polypeptide-like), deaminates cytosine and 5mC and to a lesser extent 5hmC.

**Figure 2.**
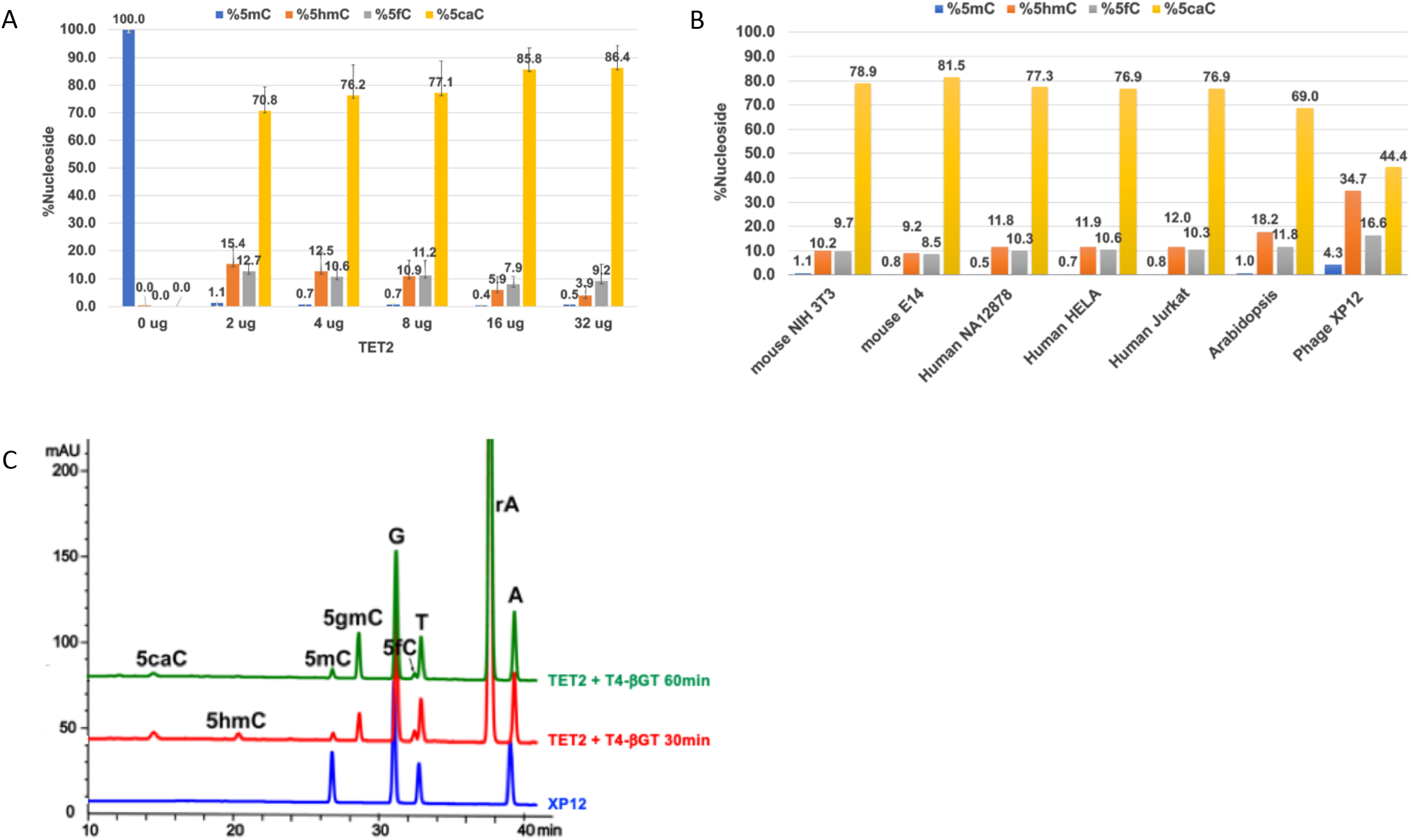
Mass Spectrometry Analysis of the Activity of TET2 and T4-βGT on 5mC and 5hmC. (A) Titration of TET2 (0 - 32 μg) with NA12878 gDNA shows that at all TET2 concentrations ≥ 99% of 5mC (blue) is oxidized to 5hmC (orange), 5fC (gray), and 5caC (yellow) with 5caC constituting ~ 70-85% of total oxidized product. (B) TET2 activity on different genomic DNAs. ≥ 99% of 5mC is converted to its oxidized forms for all gDNAs in the presence of 16 μg TET2 except for Phage XP12 where 5mC conversion was >95%. All reactions were performed with 200 ng gDNA in a total volume of 50 μL at 37 °C for 60 min. (C) LC-MS spectra showing the activity of TET2 and T4-βGT on Xp12 genomic DNA. rA is adenosine and is derived from the ATP added in to the TET2 reaction. The blue trace shows the starting Xp12 gDNA substrate. The red and green traces show the formation of four products, 5hmC, 5fC, 5caC, and 5gmC, by TET2 and T4-βGT at 30 and 60 min, respectively.

**Figure 3.**
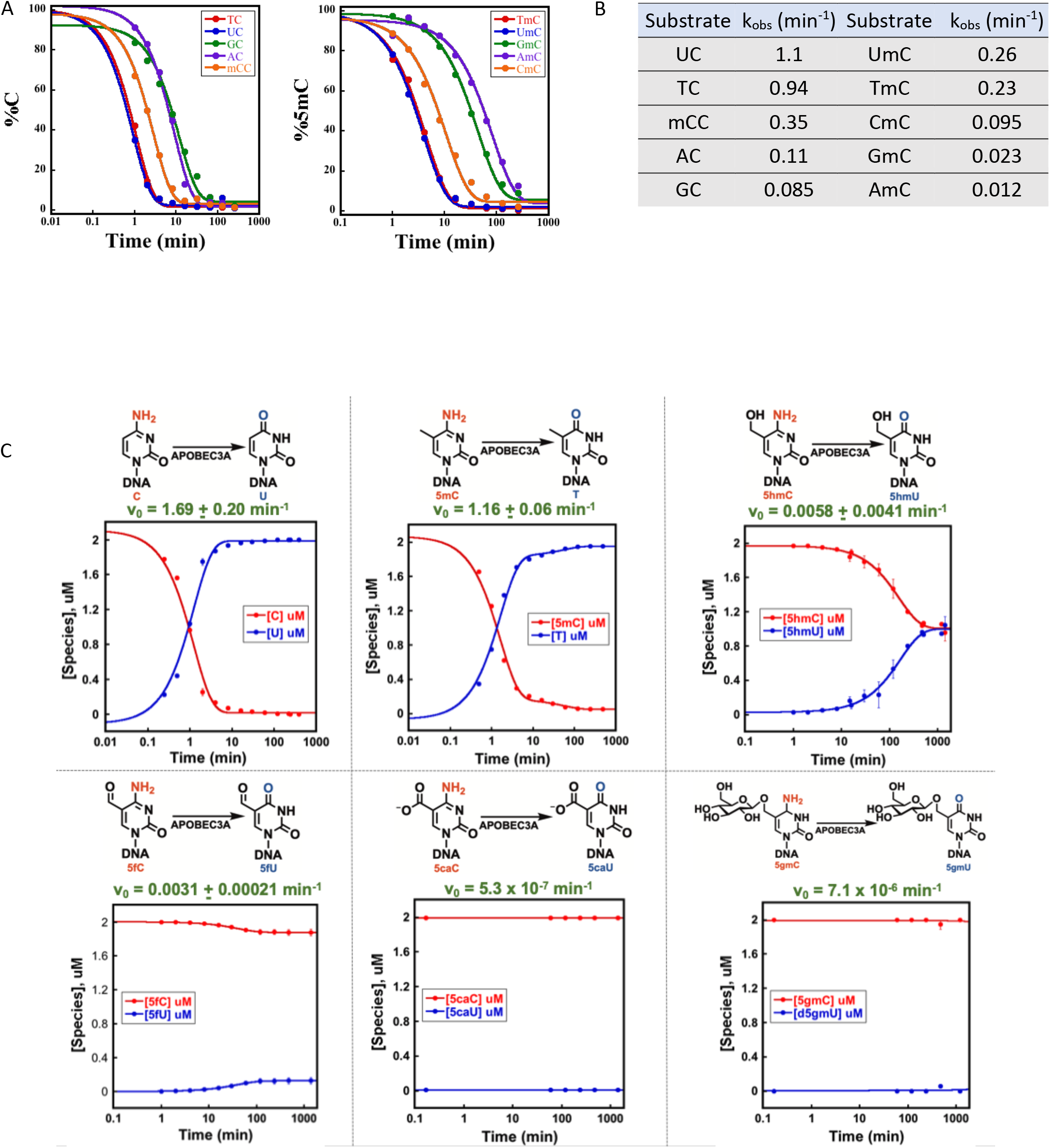
APOBEC3A/Deaminase Substrate Specificity. (A) Timecourse reactions with 0.2 μM APOBEC3A and 2 μM single-stranded oligonucleotide substrate containing either a cytosine or 5mC preceded by different bases as specified in the legend. Global nucleoside content analysis was performed by LC-MS. Oligonucleotide sequences are in Supplementary Table 1. (B) Observed rate constants (k_obs_) for C and 5mC disappearance extracted from single exponential fits (solid traces) to the experimental data (closed circles). (C) Timecourse reactions of 0.2 μM APOBEC3A and 2 μM single-stranded oligonucleotides containing either cytosine, 5mC, 5hmC, 5gmC, 5fC, 5caC (Supplementary Table 2). Synthesis of 5gmC is described in Supplementary Figure 2. k_obs_ and initial rate values for the disappearance of the various C forms are extracted from single exponential fits (solid traces) to the experimental data (closed circles) as detailed in Materials and Methods. LC-MS was used to determine the amounts of substrate and products within each reaction and the data points underwent best fit analysis using KaleidaGraph (Synergy Software).

### Activity of TET2 and T4-βGT on 5mC and 5hmC

A previously described engineered form of mouse TET2, mTET2CDΔ (16) was used in the experiments described throughout this paper and will be referred to as TET2. TET2 oxidizes greater than 99.5% of 5mC sites in NA12878 genomic DNA (gDNA) with up to ~85% being found in the 5caC oxidized form (Figure 2A). The oxidation efficiency of TET2 on 5mC was examined in gDNA from different organisms (Figure 2B) with 99% and higher of the 5mCs being efficiently oxidized in mouse and human gDNAs as well as *Arabidopsis thaliana*, which has methylation occurring in CpG (24%), CHG (6.7%), and CHH (1.7%) contexts (17). Furthermore, TET2 oxidizes 96% of 5mC on *Xanthomonas oryzae* (bacteriophage XP12) gDNA (Figure 2B). The 3% difference in the efficiency for the oxidation of XP12 gDNA compared to mammalian or Arabidopsis gDNA is attributed to XP12 having 100% of its cytosines methylated and therefore it is a far more difficult substrate for TET2 than mammalian or plant gDNA (for example only 1.5% methylation of cytosine content is found in human gDNA (18)). In addition to TET2, T4-βGT is an important component of the enzymatic system that enables discrimination of 5mC/5hmC from unmodified cytosine by glucosylating 5hmC and blocking APOBEC3A deamination. The results obtained show that T4-βGT effectively glucosylates all 5hmC formed by TET2 to 5gmC. 5fC and 5caC are also formed during the course of the TET2 reaction (Figure 2A). The combined action of TET2 and T4-βGT ensures that 5mCs and 5hmCs are modified (Figure 2C) and are no longer substrates for APOBEC3A.

### APOBEC3A substrate specificity

The deamination activity of an engineered form of APOBEC3A (NEB E7133, Ipswich, MA) was determined using single stranded oligonucleotides that share the same core sequence and contain a single cytosine or methylated cytosine preceded by either thymine, uracil, guanine, adenine, or 5mC/cytosine (Supplementary Table 1). In general, APOBEC3A favored cytosine containing oligonucleotides over their 5mC counterparts by 5-10 fold depending on the dinucleotide combination (Figure 3A, 3B). Oligonucleotides with either thymine or uracil preceding cytosine or 5mC (Supplementary Table 1) showed higher observed rate constants (kobs) for deamination than the rest of the substrates (Figure 3B). However, by extending the reaction time, all substrates were fully deaminated.

In addition to APOBEC3A activity based on dinucleotide context, we tested more extensively the effect of APOBEC3A on cytosine and several of its derivatives. Time course reactions confirm that both cytosine and 5mC are deaminated efficiently by APOBEC3A with cytosine being approximately a 2-fold better substrate (Figure 3C). On the other hand, only 50% of 5hmC substrate is deaminated by the enzyme indicating that there could possibly be a product inhibition effect. 5fC is a very poor substrate with minimal deamination detected whereas 5caC and 5gmC show no reactivity with APOBEC3A. These data demonstrate that APOBEC3A selectively deaminates cytosine and 5mC and partially deaminates 5hmC while exhibits no activity on 5fC, 5caC, and 5gmC. Therefore, an enzymatic approach for detecting 5mC and 5hmC that uses TET2, T4-βGT, and APOBEC3A can be used for specific mapping of 5mC and 5hmC within the genome.

### TET2 and APOBEC3A maintain DNA integrity

Sodium bisulfite treatment of DNA is destructive. We postulated that non-chemical conversion of DNA to detect 5mC and 5hmC using the enzymes described above for oxidation, glucosylation and deamination would maintain DNA integrity. To evaluate this, bisulfite and enzymatically converted NA12878 gDNA was PCR amplified using primers that were designed to amplify amplicons ranging in size from 543 bp to 2945 bp. Enzymatically converted DNA generated PCR products that ranged in size from 543 bp to 2945 bp (Supplemental Figure 3). Bisulfite converted DNA however, could only amplify up to the 1181 bp PCR product. These data strongly indicate that DNA is more intact using enzymatic conversion compared to chemical conversion and therefore can facilitate better detection of 5mC and 5hmC across the genome.

### EM-seq is superior to bisulfite sequencing

Taking advantage of these initial studies into TET2, T4-βGT and APOBEC3A and their potential for detection of 5mC and 5hmC while maintaining DNA integrity, EM-seq, a high-throughput sequencing method was developed. It incorporates the oxidation/glucosylation (TET2/ T4-βGT) and deamination (APOBEC3A) steps with Illumina NEBNext library preparation to produce a method that is robust, accurate and reproducible in the characterization of CpG modifications (Figure 4). EM-seq and bisulfite libraries were made using 10, 50 and 200 ng of NA12878 gDNA. Compared to bisulfite libraries, EM-seq libraries produce higher library yields using fewer PCR cycles for all DNA inputs (Figure 5A) and they also contained fewer sequencing duplicates resulting in more useable reads (Figure 5B). This ultimately increases the effective coverage of the EM-seq libraries across the genome. In addition, EM-seq libraries display a more normalized GC bias profile than bisulfite libraries which have an AT rich and GC poor profile (Figure 5C). Combining this observation with its even dinucleotide profile, (Supplementary Figure 4) provides compelling evidence that EM-seq libraries do not contain the same biases of bisulfite sequencing. These basic metrics provide the basis for the subsequent CpG analysis and the superior performance demonstrated by the EM-seq libraries.

**Figure 4.**
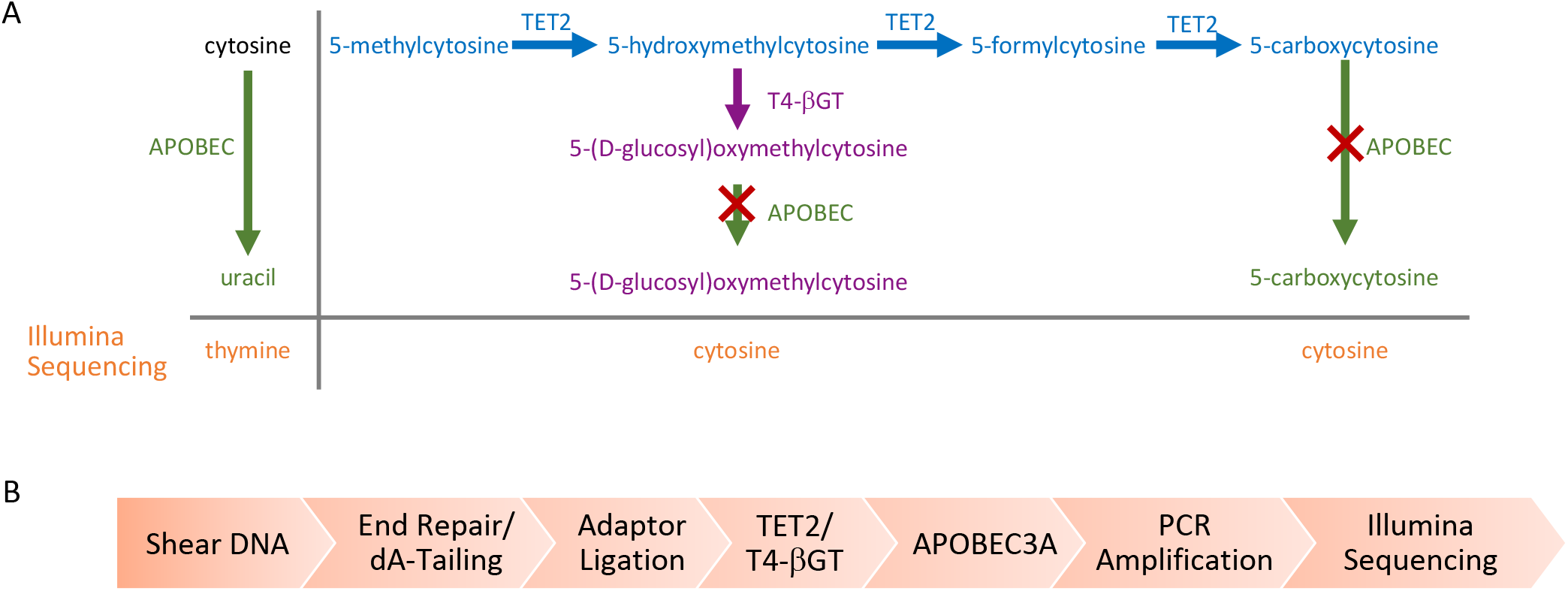
Enzymatic Methyl-seq mechanism of action and workflow. (A) Principle pathways that are important for enzymatic identification of 5mC and 5hmC using Enzymatic Methyl-seq. The actions of TET2 (blue) and T4-βGT (purple) on 5mC and its oxidation products, as well as the activity of APOBEC3A (green) on cytosine, 5gmC and 5caC are shown. The red cross represents no APOBEC3A activity. T4-βGT glucosylates 5hmC (pre-existing 5hmC and that formed by the action of TET2). TET2 converts 5mC through the intermediates 5hmC and 5fC into 5caC. APOBEC3A has limited activity on 5fC and undetectable activity on 5gmC and 5caC (Figure 3C). Uracil is replaced by thymine during PCR and is read as thymine during Illumina sequencing. (B) DNA is sheared to approximately 300 bp, end repaired and 3’ A-tailed. EM-seq adaptors are then ligated to the DNA. The DNA is treated with TET2 and T4-βGT before moving to the deamination reaction. The library is PCR amplified using EM-seq adaptor primers and can be sequenced on any Illumina sequencer.

**Figure 5.**
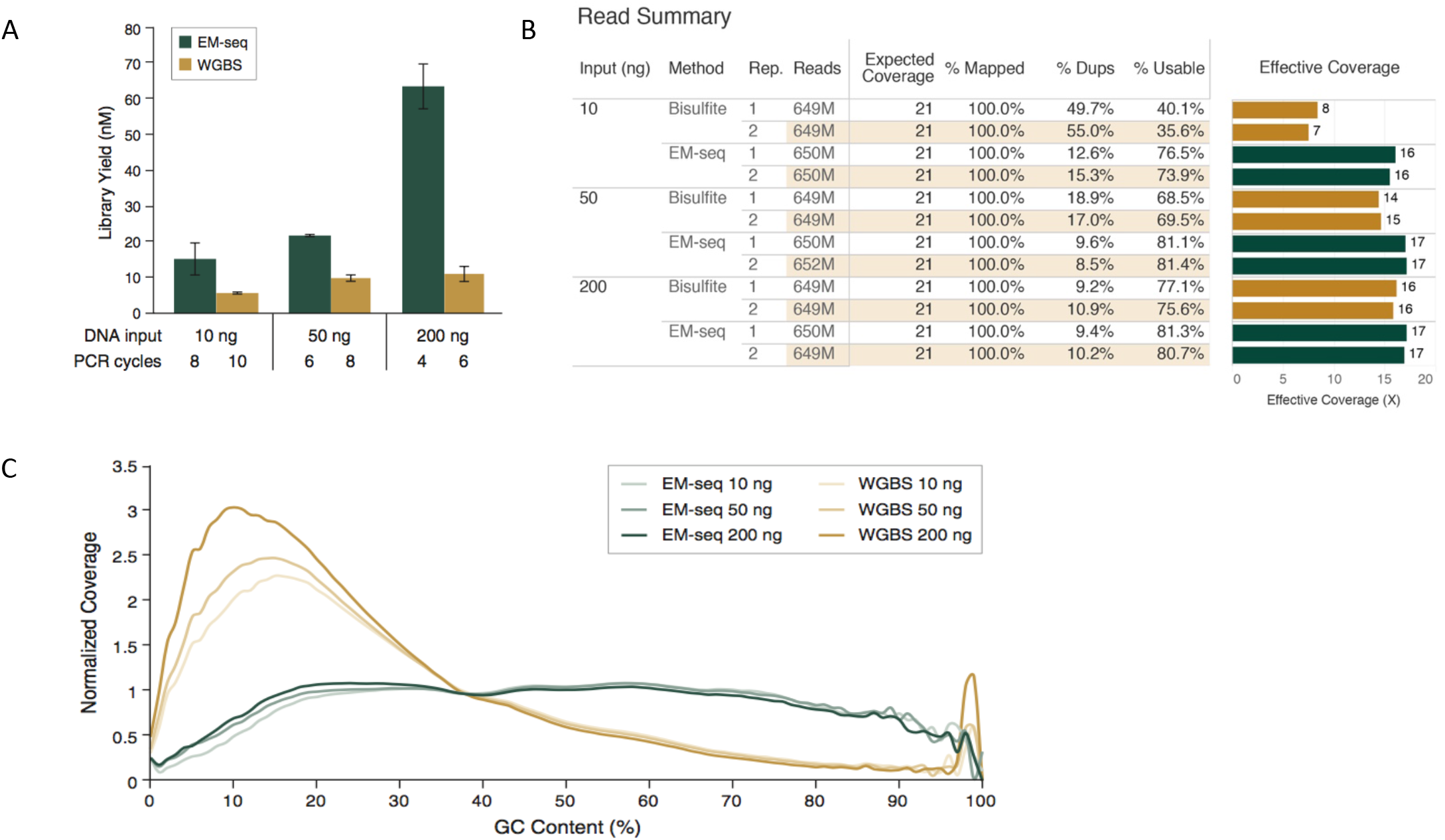
EM-seq Illumina libraries are superior to bisulfite libraries. 10 ng, 50 ng or 200 ng of NA12878 DNA were spiked with control DNA (2 ng unmethylated lambda DNA and 0.1 ng CpG methylated pUC19) and Illumina libraries were made using either EM-seq or the Zymo Gold Bisulfite Kit prepped for Illumina sequencing using NEBNext Ultra II DNA library kit reagents. Libraries were sequenced on an Illumina NovaSeq 6000 and 324 M read pairs per library were used for analysis. (A) EM-seq uses less PCR cycles but results in more PCR product than whole genome bisulfite libraries (WGBS) for all NA12878 input amounts. (B) Table of sequencing and alignment metrics for EM-seq and WGBS libraries using 324 million Illumina read pairs. Metrics were calculated using bwa-meth, Samtools, and Picard. Theoretical coverage is calculated using the number of bases sequenced/total bases in the GRCh38 reference. Percent mapped refers to reads aligned to the reference genome (grch38+controls), Percent Dups refers to reads marked as duplicate by Picard MarkDuplicates, Percent Usable refers to the set of Proper-pair, MapQ 10+, Primary, non-Duplicate reads used in methylation calling (samtools view -F 0xF00 -q 10). Effective Coverage is the Percent Usable * Theoretical coverage. (C) GC-bias plot for EM-seq and WGBS libraries. EM-seq libraries display an even GC distribution while WGBS libraries have an AT-rich and GC-poor profile.

EM-seq and bisulfite NA12878 libraries have similar cytosine methylation for all DNA inputs: CpG methylation is around 52% while for CHG and CHH contexts methylation is <0.6% (Figure 6A). Two internal controls, unmethylated lambda and CpG methylated pUC19, were included during library construction. Cytosine methylation of unmethylated lambda DNA in CpG, CHG and CHH contexts were <0.6% and for pUC19, CpG methylation was around 96% and <1.6% in the CHG and CHH contexts (Supplemental Figure 5). Furthermore, CpG coverage and the percent CpG methylation per base are represented by data analyzed using methylKit (Supplemental Figure 6). These data also demonstrate increased coverage of CpGs by EM-seq as well as the expected percentage of methylation per base with the majority of CpGs being found at 0% and 100%. Intermediate methylation is also more apparent with EM-seq than bisulfite sequencing. We also compared the methylation state between EM-seq and bisulfite libraries at increasing coverage depths. There are 56 million CpGs in the human genome, considering both DNA strands [1]. EM-seq detects approximately 54 million CpGs for the NA12878 genome at a coverage depth of 1x for the 10, 50 and 200 ng inputs, however, bisulfite libraries cover at best approximately 45 million CpGs for 50ng and 200ng inputs, but only 36 million CpGs for the 10ng input (Figure 6B and 6C). Furthermore, when the sequencing depth is increased to 8x, CpGs covered by bisulfite drop significantly compared to EM-seq. EM-seq libraries covered approximately 7 times more CpGs at 10ng inputs and 2.2 times more CpGs at 200ng inputs (Figure 6B and 6C). However, at the highest coverage depths >11x, bisulfite covers many more CpGs (Figure 6B). This is a result of the biased nature of the bisulfite libraries showing excess coverage over AT rich regions and uneven genome coverage (Figure 5B, Supplemental Figure 5). In order to compare the number of CpGs covered by EM-seq and bisulfite sequencing, correlation analysis was performed at a 1x coverage depth. Correlations between only EM-seq libraries across input amounts were consistently better compared to bisulfite libraries (Figure 6D and 6E). Expansion of the correlations to include comparison of EM-seq and bisulfite libraries (Supplemental Figure 7) also demonstrates similar patterns of correlations, with EM-seq outperforming bisulfite libraries. Overall this data indicates that EM-seq does identify the same CpGs as bisulfite as well as other CpGs that are only identified using EM-seq.

**Figure 6.**
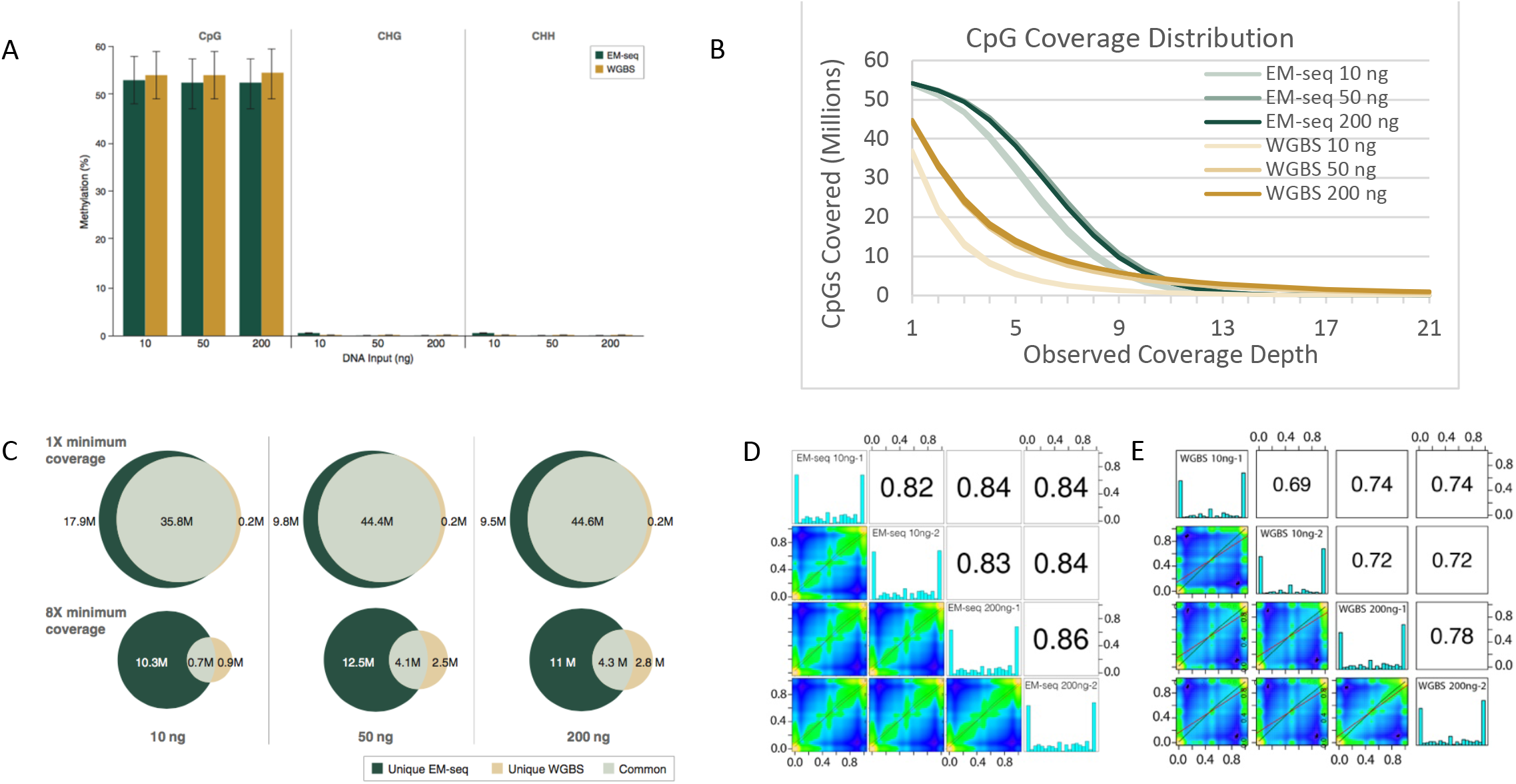
EM-seq accurately represents methylation. EM-seq and bisulfite libraries were made using 10, 50 and 200 ng of NA12878 DNA with control DNA (2 ng unmethylated lambda and 0.1 ng CpG methylated pUC19). Libraries were sequenced on an Illumina NovaSeq 6000 (2 x 100 bases). 324 million paired reads for each library were aligned to a human + control reference genome (see supplemental materials) using bwa-meth 0.2.2. and methylation information was extracted from the alignments using MethylDackel. The top and bottom strand CpGs were counted independently, yielding a maximum of 56 million possible CpG sites. (A) NA12878 EM-seq and whole genome bisulfite library (WGBS) methylation in CpG, CHH and CHG contexts are similarly represented. Methylation state for unmethylated lambda control and CpG methylated pUC19 control DNAs are shown in Supplemental Figure 5. (B) The number of CpGs covered for EM-seq and bisulfite libraries were calculated and graphed at minimum coverage depths of 1x through 21x. (C) The number of CpGs detected were compared between EM-seq and bisulfite libraries at 1x and 8x coverage depths. CpGs unique to EM-seq libraries, bisulfite libraries or those that were common to both are represented in the Venn diagrams. (D, E) Methylkit analysis at minimum 1x coverage shows good CpG methylation correlation between 10 ng and 200 ng NA12878 EM-seq libraries (D) and WGBS libraries (E). Methylation level correlations between inputs and replicates of EM-seq libraries are better than for WGBS libraries. The reduction in observations of disagreement (upper left and lower right corners) is particularly striking. Correlation between EM-seq and WGBS libraries at 10 ng, 50 ng, and 200 ng NA12878 DNA input are shown in Supplemental Figure 7.

Increased CpG coverage in EM-seq libraries also translates into an increased number of CpGs identified in genomic features (Figure 7). Interestingly, there is minimal shift in the pattern of genomic feature representation as input amount varies between the 10 to 200 ng NA12878 gDNA inputs (Figure 7A). The Dfam 3.1 list of repetitive DNA elements shows more even coverage (Figure 7B). Additional analysis of CpG coverage and their associated methylation within 2 kb of the TSS at 1x coverage depth again shows that EM-seq libraries have a higher depth of coverage compared to bisulfite libraries for all three DNA inputs (Figure 7C). Furthermore, requiring a coverage depth to 8x demonstrates not only higher coverage (Figure 7D), but that the EM-seq libraries more confidently represent the cytosine methylation status (Figure 7E). Transcription start sites are expected to have limited CpG methylation. EM-seq libraries display low levels of CpG methylation across the TSS. These results are not confined to TSS. Coverage data for other genomic features and regulatory elements is similarly even (Supplemental Figure 8).

**Figure 7.**
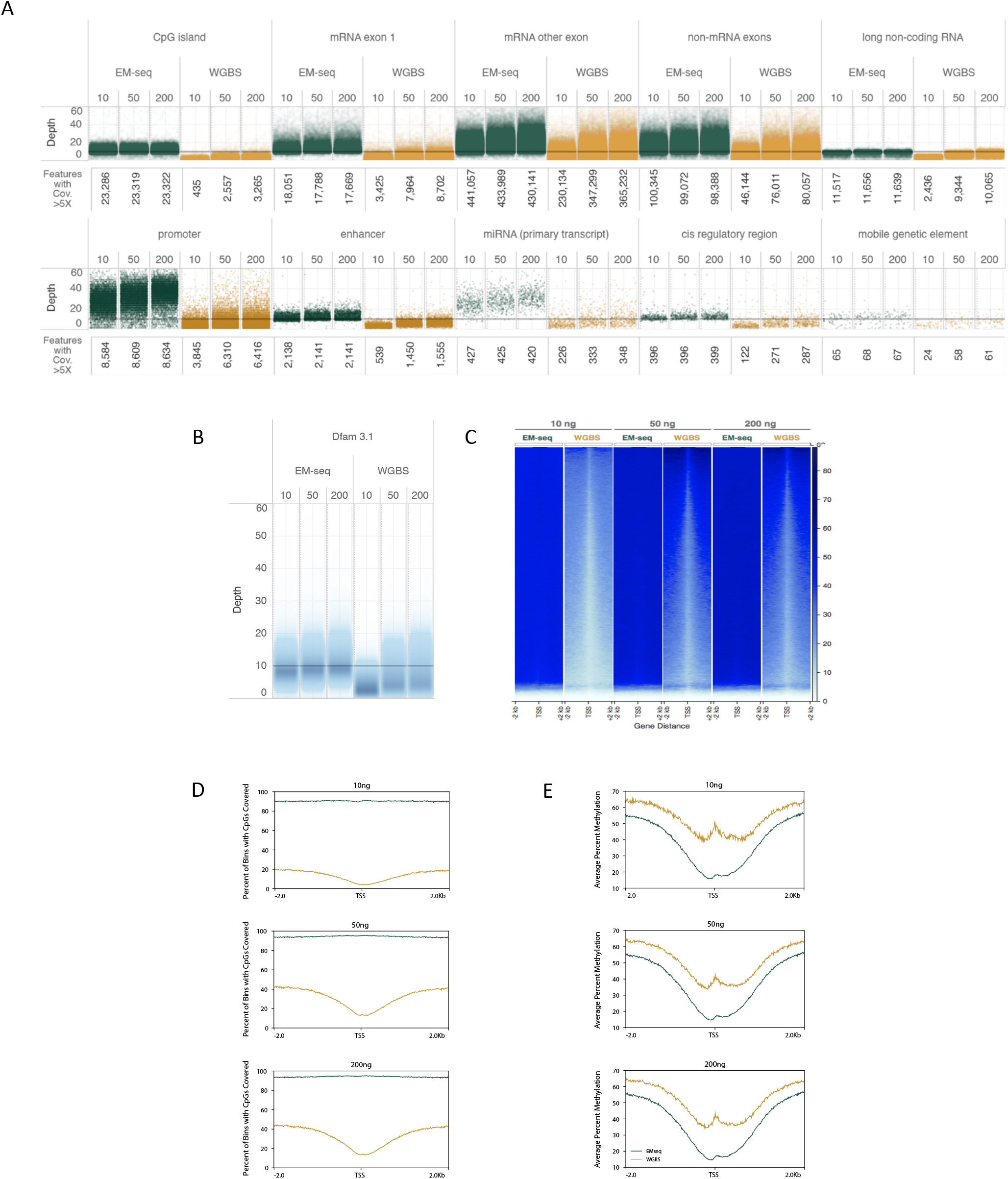
EM-seq Accurately determines methylation status of cytosines at key genomic features. 324 million EM-seq and bisulfite paired reads were analyzed for each library and annotated using featureCounts. (A) EM-seq and WGBS cover a diverse range of genomic features but EM-seq libraries exhibit greater coverage of all features examined. Coverage of various genomic feature types are represented with one point per region. The vertical position is defined by the average coverage of the feature. Points are staggered horizontally to avoid too much overlapping. Features from NCBI’s RefSeq, the Eukaryotic Promoter, UCSC table browser and Dfam are shown and the numbers covered at 5x or greater depth are indicated. (B) Repetive genomic regions (Dfam 3.1) are more evenly covered by EM-seq libraries. (C) Heat map of CpG coverage for +/- 2kb of the TSS for the three DNA inputs for EM-seq and bisulfite libraries at a minimum coverage of 1x. (D) Percent of CpGs covered at a minimum of 8x coverage, +/- 2kb, across transcription start sites (TSS) and (E) Methylation status for EM-seq and bisulfite libraries using 8x minimum coverage depth. EM-seq libraries show less CpG methylation and more accurately represent the expected CpG methylation pattern.

To further investigate whether EM-seq can be used to generate libraries using DNA inputs less than 10 ng, libraries were made using 100 pg to 10 ng NA12878 gDNA. 810 million total human reads were sampled from low input libraries and relevant standard protocol libraries. Sequencing metrics are shown in Supplemental Figure 9. These low input libraries displayed the same characteristics observed for 10 - 200 ng DNA libraries. Global methylation data for these libraries in the CpG, CHG and CHH contexts (Figure 8A, Supplemental Figure 10) were similar to standard EM-seq libraries and all libraries had even coverage across GC bias plots (Figure 8B). Due to additional PCR cycles, the number of unique reads were reduced in the 100 pg and 500 pg libraries. This is reflected in the minimum coverage plot where at 1x minimum coverage depth, approximately 24 million and 50 million CpGs were detected respectively. It is worthwhile noting that for the 10 ng WGBS libraries, only 37 million CpGs were detected, further demonstrating the increased ability of the EM-seq method to identify CpG methylation state over WGBS. The 1 ng and 10 ng low input EM-seq libraries both covered 54 million CpGs, the same number as identified using the standard protocol 10-200 ng libraries. Further analysis of CpG methylation using correlation plots (Supplemental Figure 11), comparison of the number of genomic features observed (Supplemental Figure 12), as well as heatmaps showing CpG coverage over specific genomic features (Supplemental Figure 15) all indicate that the low-input libraries perform very well. The 500 pg – 10 ng DNA inputs identified similar numbers of genomic features at >5x coverage depths and the 100 pg input covered slightly fewer (Figure 8C, Supplemental Figure 12A), likely due to additional PCR duplication. In addition, methylKit CpG methylation plots indicate that methylation is as expected with the majority of CpGs falling into either 0% methylation or 100% methylation (Supplemental Figure 13). In order to further compare the low input EM-seq method to standard EM-seq libraries and WGBS libraries, 10 ng inputs for both library types were downsampled to 810 million total human reads. In all comparisons, from correlation (Supplemental Figure 11B), methylation distribution histograms (Supplemental Figure 14) and genomic feature coverage (Supplemental Figures 12B, 15D, 15E) show that the low input and standard EM-seq methods produced similar results, with higher number of CpGs or genomic features detected and more even coverage when compared to WGBS libraries.

**Figure 8.**
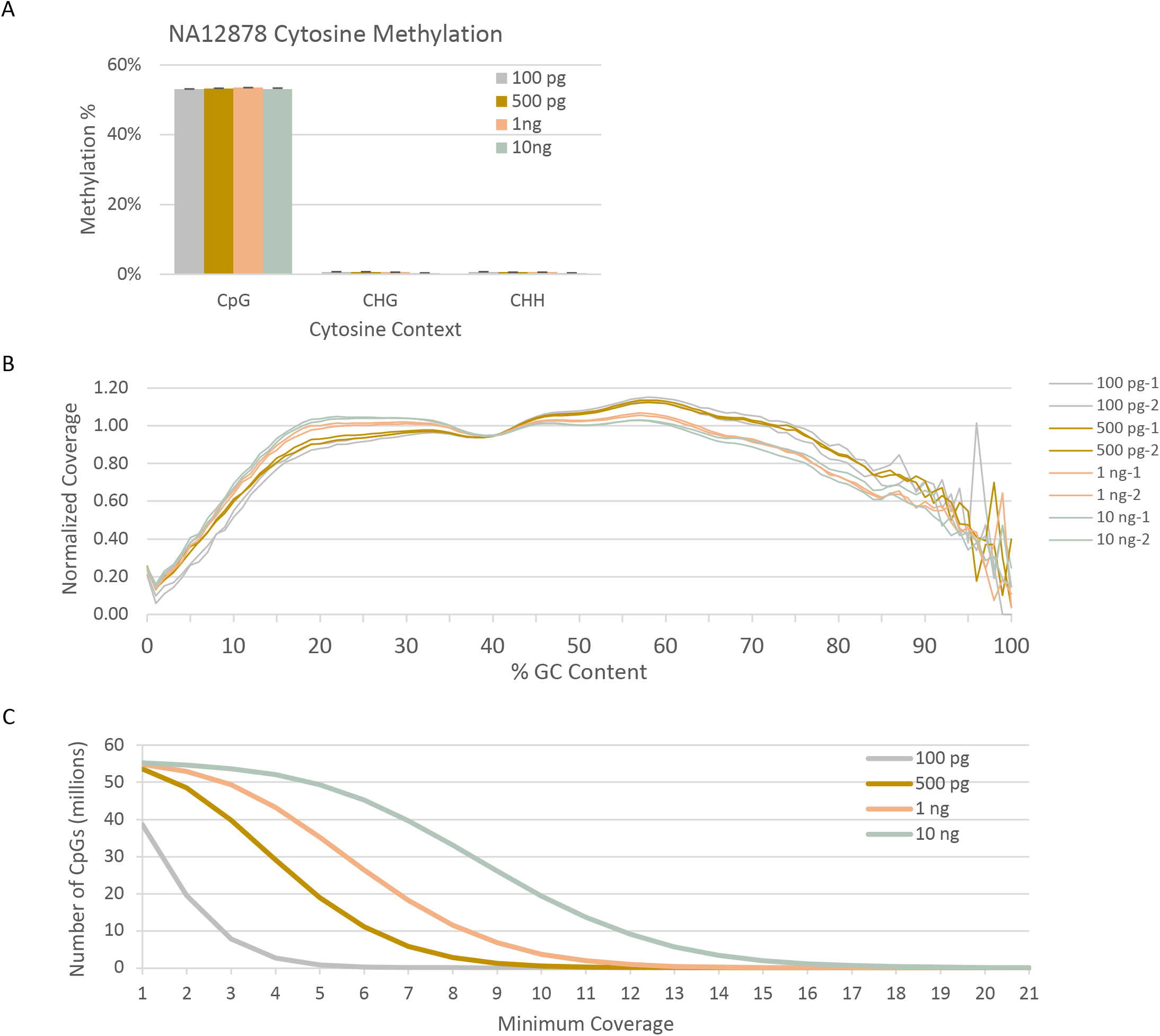
Low DNA input EM-seq libraries. Low input EM-seq libraries were made using 100 pg, 500 pg, 1 ng and 10 ng of NA12878 DNA. Libraries were sequenced on a NovaSeq 6000 and evaluated for consistency before combining technical replicates and sampling 810 million total human reads from each for methylation calling. (A) Methylation levels are as expected for all DNA inputs at around 53%. (B) GC-bias plot for EM-seq library replicates show even GC distribution. (C) CpG coverage as a function of minimum coverage depth is similar to standard-protocol libraries and shows the impact of library complexity and duplication on coverage.

## Discussion

The most routinely applied method to detect 5mC and 5hmC in gDNA uses sodium bisulfite. Despite its popularity, it does have drawbacks as it is known to damage DNA and is often associated with under- or over-conversion of cytosines (19–21). To overcome these issues, we developed EM-seq, an enzymatic method to determine the methylation status of cytosines. This method relies on a trio of enzymes, and their ability to optimally oxidize 5mC, glucosylate 5hmC, and deaminate unmodified cytosines to thymines. The TET2 enzyme used in this study has robust activity converting 99.5% of 5mC to its oxidative forms. Combining T4-βGT with TET2 effectively protects both 5mC and 5hmC, but not cytosines, from subsequent deamination by APOBEC3A. This enzymatic manipulation of cytosine, 5mC and 5hmC enables discrimination of 5mC/5hmC from cytosine in high-throughput sequence data. 5mC and 5hmC are sequenced as cytosines whereas cytosines are sequenced as thymines. These nucleotide classifications use the same designations associated with bisulfite sequencing which allows the EM-seq method to seamlessly integrate with any preferred pipeline for bisulfite data including, Bismark (22) and bwa-meth (23). This is useful as it avoids the need to invest in the design of new analysis methods.

Analysis of EM-seq and bisulfite libraries shows that both methods have very similar global methylation, but differences become noticeable with more locus-specific analysis. Traditional bisulfite sequencing fragments DNA, making it hard to sequence all mappable CpGs in a genome whereas enzymatically converted DNA does not suffer from the same fragmentation bias. More even genome coverage is especially relevant to assessment of CpG methylation state. An important consideration in generating methylomes is the amount of sequencing required to adequately cover all the relevant sites in genome adequately. Using the same number of reads, EM-seq libraries cover more genomic features at increased depths making it more economical than bisulfite libraries. EM-seq libraries can be longer (up to 350 bp, not shown), have even GC bias profiles, have dinucleotide distributions similar to unconverted DNA libraries, and can be used to generate long amplicons. Correlation analysis of CpG methylation illustrates the reproducibility and superiority of EM-seq (Figure 6D) over bisulfite sequencing (Figure 6E). It is becoming increasingly clear that methylation plays important roles in cancer and embryonic development. Bisulfite induced damage and uneven coverage have limited methylation assessment in these and other areas of research. EM-seq’s increased coverage of CpGs, combined with the accuracy of methylation calls and flexible input requirements opens up areas of research that were previously inaccessible.

The adaptation of enzymatic conversion for high-throughput sequencing enhances the study of the whole methylome and permits DNA profiling from challenging samples. These include lower DNA inputs as well as more diverse types of input material. Bisulfite sequencing has traditionally been difficult to apply to cell free DNA and Formalin Fixed Paraffin Embedded (FFPE) DNA, primarily due to low inputs and the presence of DNA damage. The EM-seq results herein demonstrate that the method can overcome these challenges and provides researchers with the required tools to probe new avenues of research. We demonstrated that EM-seq can be used with as little as 100 pg DNA and that the methylation and CpG coverages follow similar trends as seen with 10-200 ng DNA inputs. In addition to short-read sequencing, enzymatic conversion has the potential to be optimized for long read sequencing using PacBio or Oxford Nanopore, as well as enzymatic equivalents for reduced representation bisulfite libraries (RRBS), use of converted DNA to probe arrays, perform target enrichment and long-amplicon sequencing. We envision most of the improvements to these applications coming from the intact nature of the DNA that is associated with the enzymatic conversion found in EM-seq and resultant improvements to coverage profiles.

New methods to capture the methylation or hydroxymethylation status of cytosines have recently been reported. TET-assisted pyridine borane sequencing (TAPS) combines enzymatic oxidation of 5mC to 5caC with a chemical deamination step of 5caC. The method can also be modified to detect 5hmC, by glucosylating 5hmC to 5gmC prior to TET oxidation. Another method, ACE-seq relies solely on enzymes to glucosylate 5hmC before deaminating both cytosine and 5mC to uracil but provides no information on 5mC. Both of these methods have drawbacks. TAPS requires significant investment in the development and validation of new analysis tools for data analysis, shows more biased coverage of CpG Islands than EM-seq, and has not been shown to work with inputs below 1 ng of DNA (15). Sequencing data produced from EM-seq libraries can be analyzed using pipelines currently used for bisulfite data, shows almost no coverage bias toward any specific genomic feature and can work with inputs as low as 100 pg of DNA. Combining ACE-seq data with bisulfite data can identify 5mCs, but this method is limited by the quality of the bisulfite data. EM-seq detects both 5mC and 5hmC and the method can be modified to only detect 5hmC, in a similar manner to that of ACE-seq to produce a more complete picture of the methylome.

In stark contrast to bisulfite converted sequencing libraries, the DNA incorporated into EM-seq libraries suffers from little to no damage and has very few biases associated with it. This combined with the fact that the EM-seq method enables accurate calling of DNA methylation, enhances the potential of this method for extraction of information from challenging samples, including low input, FFPE damaged and cell free DNA, and therefore enabling discovery in areas of methylome research that have been inaccessible to bisulfite sequencing.

Many researchers are pursuing multi-omics based approaches to better elucidate the various mechanisms involved in gene regulation. Inclusion of EM-seq methylome information in these studies will provide a more complete understanding of epigenomics and therefore when combined with mRNA expression levels (RNA-seq), chromatin state determinations (ATAC-seq, NOMe-seq, NicE-seq, Hi-C, Histone modifications), and regulatory factor occupancy (ChiP-seq) data would lead to enhanced multi-omic comparisons. This ultimately will lead to a more complete understanding of cell regulation.

## Conclusions

EM-seq provides accurate characterization of cytosine methylation within genomes. Comparison of data generated using EM-seq to that of bisulfite sequencing, the current gold standard, demonstrates that EM-seq is a vastly superior method. EM-seq does not damage DNA in the same ways as reported for bisulfite sequencing. Subsequently, all metrics examined, including but not limited to genome coverage, CpGs detected and genomic features identified, are improved compared to bisulfite sequencing. The ability to use lower DNA inputs with EM-seq in combination with retaining essential metrics that are typically lost using bisulfite sequencing shows the versatility of this method. EM-seq can also be used in other applications requiring low DNA input or damaged DNA such as cell free DNA, single cells or FFPE DNA. These types of DNA input are often associated with researching development or in diagnosis of cancer.

## Materials and Methods

### TET2 and APOBEC3A protein

Generation, expression and purification of the TET2 construct mTET2CDΔ are described previously (16). The APOBEC3A protein was produced by (NEB E7133, Ipswich, MA).

### E14 Embryonic Stem (ES) Cell culture

ES cells were cultured as described in (24). Briefly, cells were grown in GMEM (Invitrogen) media containing 10% FBS (Gemcell), 1% non-essential amino acids (NEAA) (Hyclone), 1% sodium pyruvate (Invitrogen), 50 μM 2-mercaptoethanol (Sigma), and 1× LIF (Millipore). To maintain the undifferentiated state, ES cells were grown on 0.1% gelatin-coated culture dishes (Stem Cell Technologies).

### DNA substrates

Mouse NIH/3T3 and human Jurkat DNA were from NEB (Ipswich, MA), and XP12 phage DNA was obtained from Dr. Peter Weigele (NEB, Ipswich, MA). E14 gDNA was extracted using QIAGEN DNeasy Blood and Tissue Kit following manufacturer’s instructions (QIAGEN). NA12878 gDNA was obtained from Corriell Cell Repositories (Camden, NJ). Unmethylated lambda DNA was obtained from Promega (Promega, Madison, WI) and Arabidopsis gDNA from Biochain, CA. CpG methylated pUC19, was made by transforming pUC19 plasmid (NEB, Ipswich, MA) into dam-/dcm-competent *E.coli* cells (NEB, Ipswich, MA). Unmethylated pUC19 was extracted using the Monarch Plasmid Miniprep Kit (NEB, Ipswich, MA). 10 μg of pUC19 was CpG methylated *in vitro* using 40 units of M.SssI (NEB, Ipswich, MA) in a 200 μl final reaction volume for 2 h at 37 °C. The reaction was cleaned up using NEBNext Sample Purification beads with two 80% ethanol washes. CpG methylated pUC19 DNA was eluted in 100 μl H2O before repeating the methylation reaction to minimize the presence of any unmethylated CpGs.

### Enzymes and oligonucleotide substrates

All enzymes were from New England Biolabs (Ipswich, MA) unless otherwise noted. Oligonucleotides were synthesized by IDT (Coralville, IA).

### Preparation of gDNA substrate mTET2 kinetics experiments

6 μg of gDNA was made up to 100 μl using 10mM Tris 1mM EDTA, pH 8.0 buffer and sheared to 1500 bp using the Covaris S2 instrument. The sample was sheared using settings of 5% duty cycle, intensity 3 and 200 cycles/burst for 40 s. Sheared DNA was purified using either the Monarch PCR and DNA Cleanup kit (NEB, Ipswich, MA) or QIAquick Nucleotide Removal Kit (QIAGEN). The purified DNA was eluted in H_2_O and used in the TET reaction.

### TET2 end timepoint reaction

A master mix containing 50 mM TRIS pH 8.0, 2 mM ATP, 1 mM DTT, 5 mM sodium ascorbate, 5 mM αKG (alpha-ketoglutarate), varying concentrations of TET2 and 2 ng/μl gDNA was prepared in a final volume of 50 μl. The reaction was initiated with the addition of 50 μM FeSO_4_ and incubated at 37 °C for 60 min. The DNA sample was then treated with 0.8 Units of Proteinase K (NEB, Ipswich, MA) for 60 min at 50 °C and subsequently purified using Oligo Clean & Concentrator kit (Zymo Research) and analyzed for oxidized 5mC modifications using liquid chromatography-mass spectrometry/mass spectrometry (LC-MS/MS) according to the procedure and instruments described previously (16).

### TET2 time-course reaction

A master mix containing 50 mM Tris pH 8.0, 2 mM ATP, 1 mM DTT, 5 mM sodium ascorbate, 5 mM α-KG, 0.58 μg/μl or 10 μM TET2, and 50 μM FeSO_4_ was prepared in a 50 μl total volume and the reaction was initiated with the addition of 0.9 ng/μl XP12 gDNA and was incubated at 37 °C. 20 μl aliquots were removed at 0, 0.5, 1, 2, 15, 60, and 120 min, and added to 200 μl isopropanol to quench the reaction. The samples were centrifuged for 2 min at 13,000 rpm, dried using a SpeedVac vacuum concentrator and resuspended in 50 μl of H_2_O. 0.8 U of Proteinase K (NEB, Ipswich, MA) was added and incubated for 60 min at 50 °C and the DNA was purified using the Oligo Clean & Concentrator kit (Zymo). DNA was analyzed for oxidized 5mC modifications using liquid chromatography-mass spectrometry/mass spectrometry (LC-MS/MS) according to the procedure and instruments described previously (16).

### Glucosylation of 5hmC containing oligonucleotide

For the APOBEC3A substrate assay, a single stranded oligonucleotide containing 5hmC (Supplemental Table 2) was glucosylated at the 5hmC. 10 μg of substrate was incubated with 40 U of T4-βGT supplemented with 40 μM UDP-glucose in a 500 μl reaction for 16 h at 37°C.

### APOBEC3A substrate specificity and site preference analysis

The oligonucleotides used for the quantitative analysis of APOBEC3A site preference are shown in Supplemental Tables 1 and 2. Each oligonucleotide was assayed as follows. 2 μM ssDNA oligonucleotide and 0.2 μM APOBEC3A in 1x reaction buffer (50 mM Bis-Tris, pH 6.0, 0.1% Triton X-100) were incubated at 37°C. Incubation times varied according to the oligonucleotide being assayed. For the site preference analysis, timepoints of 0, 1, 2, 4, 8, 20 h were used. For substrate specificity assays, C and 5-mC incubation times were 1, 2, 4, 8, 16, 32, 64, 128, and 256 min. For the oligonucleotides containing 5hmC, 5fC or 5caC, time points of 8, 16, 32, 64, 128, 256 min, 5 and 22 h were used. Reactions were quenched with 8 volumes of ethanol and the DNA was recovered by using the DNA Clean and Concentrator kit (Zymo Research). DNA samples were digested to nucleosides using the Nucleoside Digestion Mix (NEB, Ipswich, MA). Global nucleoside content analysis was performed by LC-MS or LC-MS/MS as noted (see conditions below). For the site preference and substrate specificity assays, the data points were best fitted by a single exponential equation to follow the disappearance of the appropriate nucleotide: C, 5mC, 5hmC, 5fC, or 5caC or U and T appearance (KaleidaGraph, Synergy Software).

### Global nucleoside analysis using LC-MS and LC-MS/MS

LC-MS analysis was performed on an Agilent LC-MS System 1200 Series equipped with a G1315D diode array detector and a 6120 Single Quadrupole Mass Detector operating in positive (+ESI) and negative (-ESI) electrospray ionization modes. LC was carried out on a Waters Atlantis T3 column (4.6 × 150 mm, 3 μm) with a gradient mobile phase consisting of 10 mM aqueous ammonium acetate (pH 4.5) and methanol. The relative abundance of each nucleoside was determined by UV absorbance. LC-MS/MS analysis was performed in duplicate by injecting digested DNA on an Agilent 1290 UHPLC equipped with a G4212A diode array detector and a 6490A Triple Quadrupole Mass Detector operating in the positive electrospray ionization mode (+ESI). UHPLC was carried out on a Waters XSelect HSS T3 XP column (2.1 × 100 mm, 2.5 μm) with the gradient mobile phase consisting of 10 mM aqueous ammonium formate (pH 4.4) and methanol. MS data acquisition was performed in the dynamic multiple reaction monitoring (DMRM) mode. Each nucleoside was identified in the extracted chromatogram associated with its specific MS/MS transition: dC [M+H]^+^ at m/z 228 →112, 5mC [M+H]^+^ at m/z 242→126, 5hmC [M+H]^+^ at m/z 258→142, 5fC [M+H]^+^ at m/z 256→140 and 5caC [M+H]^+^ at m/z 294→178. External calibration curves with known amounts of the nucleosides were used to calculate their ratios within the analyzed samples.

### Long Amplicon Generation

Sodium bisulfite conversion: 200 ng of NA12878 DNA was converted using the EZ DNA Methylation-Gold Kit (ZYMO Research, Irvine, CA) according to the manufacturer’s instructions. Enzymatic 5mC/5hmC Conversion: 200 ng of NA12878 DNA was incubated with 16 μg of TET2 enzyme in a buffer containing 50 mM Tris pH 8.0, 2 mM ATP, 1 mM DTT, 5 mM sodium ascorbate, 5 mM a-KG and 50 μM Fe(II) for 30 min at 37°C, in a final volume of 50 μl. 20 U of T4-βGT (NEB, Ipswich, MA) and 1 μl of 50x UDP-glucose were added and incubated for 30 min at 37°C. Then, 0.8 U of Proteinase K (NEB, Ipswich, MA) was added and incubated for 30 min at 37°C. DNA was purified using a Genomic DNA Clean & Concentrator (ZYMO Research, Irvine, CA). The DNA was then denatured by adding formamide to 20% (v/v) at and incubating for 10 min at 90°C. DNA was deaminated using 0.26 μg of APOBEC3A in a 100 μl reaction volume for 3 h. 3 μl from 100 μl of enzymatically converted and 1 μl from 15 μl of bisulfite-converted DNA were PCR amplified with Q5U polymerase (NEB, Ipswich, MA) and primers (Supplemental Table 3) using the following cycling conditions: 98°C for 30 s, then 35 cycles of 98°C for 10 s, 63°C for 20 s, and 68°C for 2 min. The resulting PCR products were 543, 1181, 1750, 2207 and 2945 bp. PCR amplicons were visualized on a 2% agarose gel run in 1x TBE buffer and stained with ethidium bromide.

### Enzymatic Methyl-seq

10, 50 and 200 ng of NA12878 gDNA was combined with 0.1 ng CpG methylated pUC19 and 2 ng unmethylated lambda control DNA and made up to 50 μl with 10 mM Tris 0.1 mM EDTA, pH 8.0. The DNA was transferred to a Covaris microTUBE (Covaris, Woburn, MA) and sheared to 240-290 bp using the Covaris S2 instrument. DNA was sheared twice for 40 s at Duty Cycle: 10%, Intensity: 4, Cycles/Burst: 200. The 50 μl of sheared material was transferred to a PCR strip tube to begin library construction. NEBNext DNA Ultra II Reagents (NEB, Ipswich, MA) were used according to the manufacturer’s instructions for end repair, A-tailing and adaptor ligation of 0.4 μM EM-seq adaptor (A5mCA5mCT5mCTTT5mC5mC5mCTA5mCA5mCGA5mCG5mCT5mCTT5mC5mCGAT5mC*T and [Phos]GAT5mCGGAAGAG5mCA5mCA5mCGT5mCTGAA5mCT5mC5mCAGT5mCA). The ligated samples were mixed with 110 μl of resuspended NEBNext Sample Purification Beads and cleaned up according to the manufacturer’s instructions. The library was eluted in 29 μl of water. DNA was oxidized in a 50 μl reaction volume containing 50 mM Tris HCl pH 8.0, 1 mM DTT, 5 mM Sodium-L-Ascorbate, 20 mM a-KG, 2 mM ATP, 50mM Ammonium Iron (II) sulfate hexahydrate, 0.04 mM UDG (NEB, Ipswich, MA), 16 μg mTET2, 10 U T4-βGT (NEB, Ipswich, MA). The reaction was initiated by adding Fe (II) solution to a final reaction concentration of 40 μM and then incubated for 1 h at 37°C. Following this, 0.8 U of proteinase K (NEB, Ipswich, MA) was added and before incubation for 30 min at 37°C. At the end of the incubation, the DNA was purified using 90 μl of resuspended NEBNext Sample Purification Beads according to the manufacturer’s instructions. DNA was eluted in 17 μl of water and 16 μl was then transferred to a new PCR tube and denatured by addition of 4 μl of formamide (Sigma-Aldrich, St. Louis, MO) and incubation at 85°C for 10 min. The DNA was then deaminated in 50 mM Bis-Tris pH 6.0, 0.1% Triton X-100, 20 μg BSA (NEB, Ipswich, MA) using 0.2 μg of APOBEC3A. The reaction was incubated at 37°C for 3 h and the DNA was purified using 100 μl of resuspended NEBNext Sample Purification Beads according to the manufacturer’s protocol. The sample was eluted in 21 μl water and 20 μl was transferred to a new tube. 1 μM of NEBNext Unique Dual Index Primers and 25 μl NEBNext Q5U Master Mix (M0597, New England Biolabs, Ipswich, MA) were added to the DNA and amplified as follows: 98°C for 30 s, then cycled 4 (200 ng), 6 (50 ng) and 8 (10 ng) times according to DNA input, 98°C for 10s, 62°C for 30 s and 65°C for 60 s. A final extension of 65°C for 5 min and hold at 4°C. EM-seq libraries were purified using 45 μl of resuspended NEBNext Sample Purification Beads and the sample was eluted in 21 μl water and 20 μl was transferred to a new tube. Low input EM-seq libraries for 100 pg - 10 ng gDNA inputs were processed as for the 10 – 200 ng gDNA inputs and used 2U T4-βGT. Libraries were quantified using D1000HS Tape for TapeStation (Agilent) prior to sequencing on an Illumina NovaSeq instrument.

### Whole Genome Bisulfite Sequencing

10, 50 and 200 ng of NA12878 gDNA was combined with 0.1 ng CpG methylated pUC19 and 2 ng unmethylated lambda control DNA and made up to 50 μl with 10 mM Tris 0.1 mM EDTA, pH 8.0. DNA samples were sheared using the same conditions as EM-seq and processed through NEBNext Ultra II library preparation. TruSeq DNA Single indices (Illumina, San Diego, CA) were used instead of the EM-seq adaptor. Bisulfite conversion was performed using EZ DNA Methylation-Gold Kit (Zymo Research, Irvine, CA) following manufacturer’s instructions.

### DNA Analysis

Paired-end reads from WGBS and EM-seq libraries were sequenced on the same flowcells, demultiplexed using Picard’s IlluminaBasecallsToFastq 2.18.17. Fastq reads were adapter trimmed (trimadap from bwakit (25)) and aligned to a reference genome including the GRCh38 analysis set and sequences used as controls (phage lambda, puc19c, phage T4 and phage XP12) (Supplemental Table 5) using bwameth[23]. Reads were duplicate marked (samblaster (26)) before sorting (sambamba (27)). Figures were generated from the same number of reads for each library, randomly sampled (sambamba view -t 8 -s ${frac_of_largest}). Methylation amounts by contig and context were calculated using Methyldackel mbias and extracted using Methylation extract (default settings). The GC Bias plot was generated using Picard’s CollectGCBiasMetrics and Insert size distribution was created with CollectInsertSizeMetrics.

A nextflow pipeline with full analysis detail is available (28). Briefly, unique features were assembled from the Eukaryotic Promoter Database (promoters), UCSC’s table browser (CpG Islands), Dfam (repetitive elements), and NCBI’s RefSeq annotation (remainder). Coverage of these features was assessed using featureCounts.

Coverage of Transcription start sites (GENCODE v29 (29)) was calculated using deepTools (30) after combining both technical replicates and sampling 1.3B reads for each input amount and conversion method.

BigWig coverage tabulation:

~~~
find . -name ‘*combined_reps.bam’ | parallel bamCoverage -b {} -o {.}_3840_mq30.bw \
  --samFlagExclude 3840 --minMappingQuality 30
~~~

Coverage matrix construction:

~~~
computeMatrix reference-point -S *.bw -R gencode.v29.basic.annotation.gtf \
  -a 2000 -b 2000 -o 2kb_start_codon.matrix --referencePoint TSS -p max/2
~~~

Heatmap plotting:

~~~
plotHeatmap -m 2kb_start_codon.matrix --boxAroundHeatmaps no \
  --colorList azure,lightblue,cornflowerblue,royalblue,blue,mediumblue,darkblue,navy,midnightblue \
  --missingDataColor white --yMax 65 --yMin 0 --colorNumber 80 \
  --samplesLabel “10 ng EM-seq” “10 ng WGBS” “50 ng EM-seq” “50 ng WGBS” “200 ng EM-seq” “200 ng WGBS” \
  -o gencode.v29.basic.annotation_tss_cov_blue.pdf
~~~

We used a binning approach to calculate the fraction of sites covered and methylation status of sites with at least 8x coverage. Windows of +/- 2kb around each TSS were divided into 10 bp bins. We calculated the percent of bins containing at least one CpG with at least 8x coverage in comparison to all bins with at least 8x coverage. For methylation, we calculated the average methylation of CpGs with at least 8x coverage in each 10 bp bin.

We assessed the number of “usable reads” using “samtools view -F 0xF00 -q 10 -c ${bam}”

A nextflow (31) workflow containing the analysis detail is available in the supplemental materials.

Due to excess control concentration in the low-input experiment, downsampling was performed to produce equivalent numbers of human reads and replicates were assessed independently before combination for final methylation assessment. Summary fractions are expressed as fractions of human reads (instead of total reads). Otherwise, software methods remained the same as for the standard protocol libraries. Software versions: Methyldackel 0.3.0, bwameth=0.2.0, bwa=0.7.17, bwakit=0.7.15, samblaster=0.1.24, sambamba=0.6.6, picard=2.18.14.

## Declarations

### Ethics approval and consent to participate

Not Applicable

### Consent for publication

Not Applicable

### Availability of data and materials

The sequencing data generated in this manuscript can be found at https://www.ncbi.nlm.nih.gov/bioproject/PRJNA591788

### Competing interests

This research was supported by New England Biolabs, Inc. RV, VKCP, ZS, BWL, LS, SG, ND, MAC, BSS, KM, MS, JCS, HEC, ET, IRC, SP, ETD, TCE, LW and TBD performed the work as employees of New England Biolabs, Inc.

### Funding

Funding was provided by New England Biolabs Inc.

### Authors’ contributions

TBD conceived the original idea and supervised the project. SP contributed to the original idea and helped with isolating genomic DNA. LW, VKCP, RV, LS, BSS, ZS, BWL, ETD, TBD designed the experiments. RV and HEC conducted and interpreted APOBEC3A biochemical experiments. LS designed, conducted and interpreted TET2 biochemical experiments. JCS, MS, SG, ET, KM, LS expressed and purified APOBEC3A and TET2 proteins, designed and performed biochemical assays. ND and IRC performed LCMS analysis. LW, VKCP, RV, BSS, ZS, ETD optimized EM-seq protocol. BWL, VKCP, MAC, ZS performed bioinformatic analysis. LW, VKCP, RV, ZS, BWL, LS, TCE, TBD interpreted results and co-wrote the manuscript. All authors read and accepted the final version of the manuscript.

## Acknowledgements

We would like to thank Donald Comb, James Ellard, and Rich Roberts at New England Biolabs Inc. for research support and encouragement. We also thank Laurie Mazzola, Danielle Fuchs and Kristen Augulewicz from the Sequencing Core.

## Notes

https://www.ncbi.nlm.nih.gov/bioproject/PRJNA591788

### Competing Interest Statement

The authors have declared no competing interest.

