## Supplemental Figures for "EM-seq: Detection of DNA Methylation at Single Base Resolution from Picograms of DNA"

**Supplemental Figure 1**

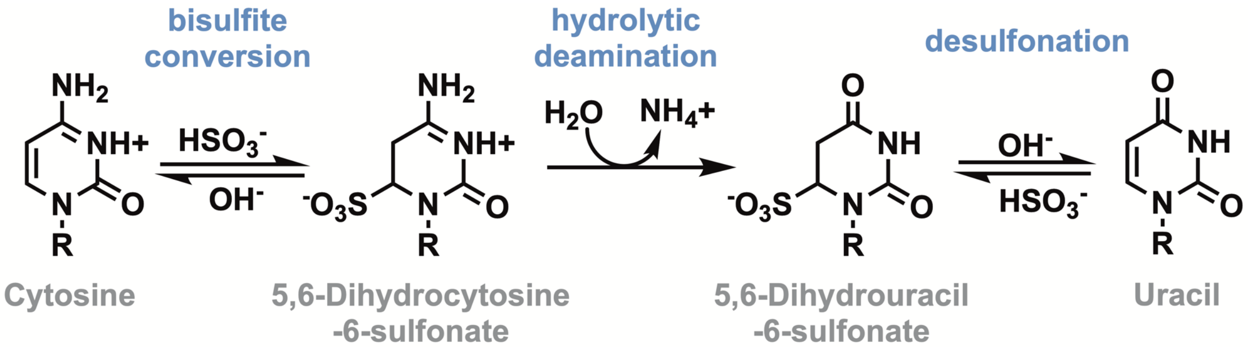


**Chemistry of sodium bisulfite conversion**

Three chemical steps are needed to deaminate cytosine to uracil. Bisulfite conversion forms the intermediate 5,6-Dihydrocytosine-6-sulfonate which can then undergo hydrolytic deamination to form 5,6-Dihydrouracil-6-sulfonate. Desulfonation during the final step of conversion results in the formation of uracil.

**Supplemental Figure 2**

A

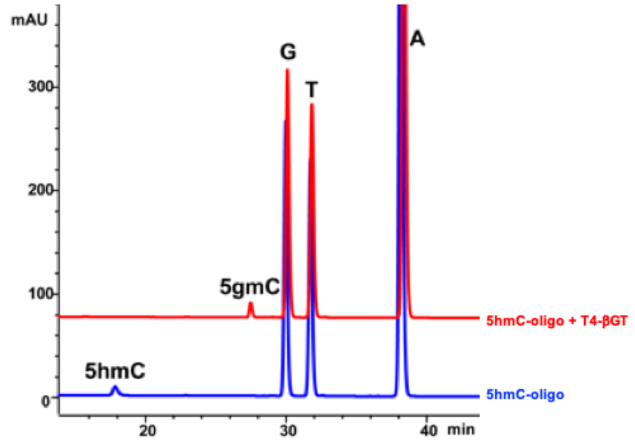


B

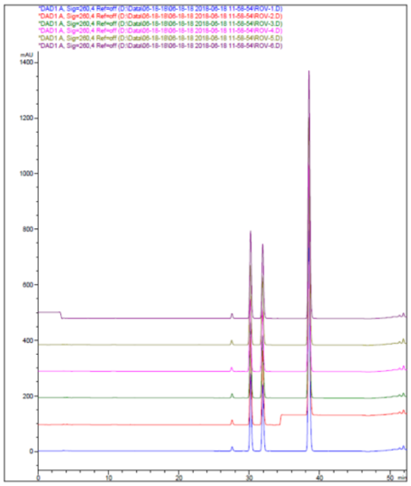

C

| APOBEC3A incubation time (h) | 5gmC | G | T | A |
|---|---|---|---|---|
| 0 | 0.036 | 0.397 | 0.507 | 1.000 |
| 1 | 0.036 | 0.399 | 0.510 | 1.000 |
| 2 | 0.036 | 0.398 | 0.509 | 1.000 |
| 4 | 0.036 | 0.397 | 0.508 | 1.000 |
| 8 | 0.035 | 0.397 | 0.507 | 1.000 |
| 20 | 0.036 | 0.398 | 0.509 | 1.000 |

**Generation of 5-(β-glucosyloxymethyl)cytosine single stranded DNA substrate**

(A) An oligonucleotide with an internal 5-hydroxymethylcytosine (ATAAGAATAGAATGAAT/i5HydMe-dC/GTGAAATGAATATGAAATGAATAGTA) was incubated with T4-βGT and the formation of 5gmC demonstrated. (B, C) Timecourse showing APOBEC3A activity on the glucosylated oligonucleotide. APOBEC3A does not deaminate 5gmC. The extinction coefficient constant for 5gmC is not available, therefore the extinction coefficient for C was used instead. This oligonucleotide was used in Figure 3C.

**Supplemental Figure 3**

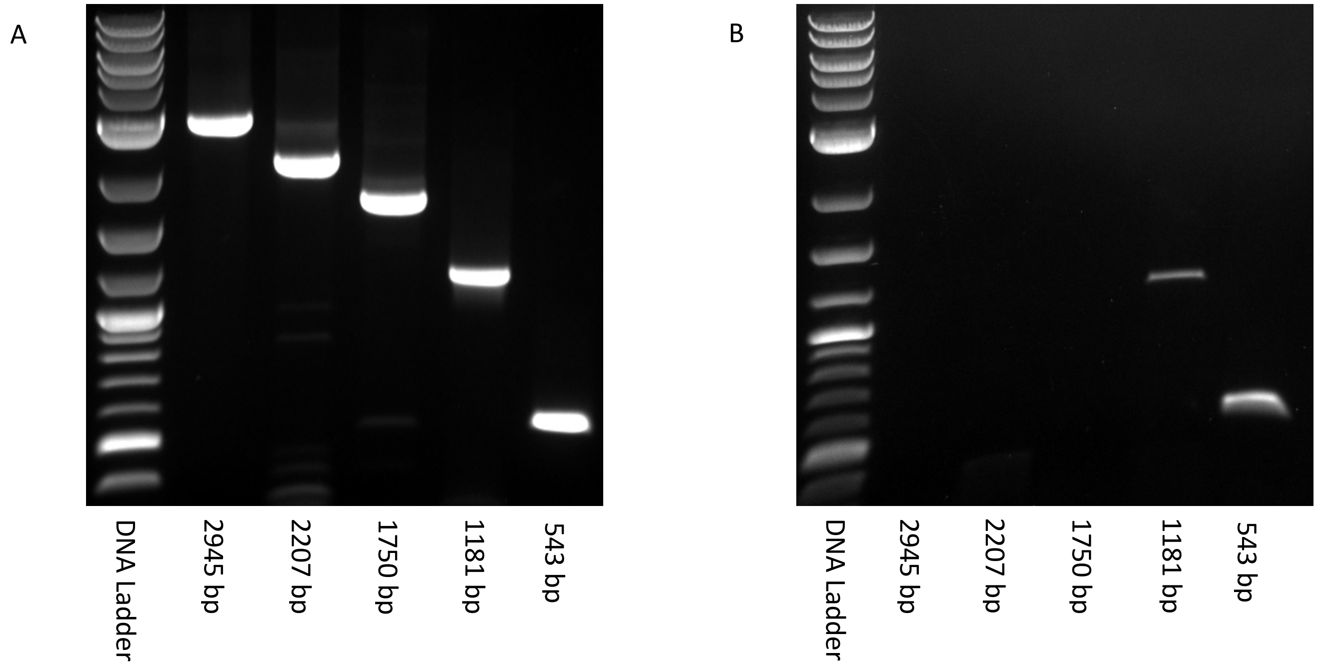


**DNA integrity assessment**

Agarose gel images of end-point PCR with five amplicons ranging in size from 543 bp - 2945 bp. NA12878 DNA was sheared to 5 kb prior to enzymatic conversion or bisulfite conversion. The PCR primers used are shown in Supplementary Table 3. (A) TET2/T4-βGT followed by APOBEC3A treated samples and (B) bisulfite treated DNA. The left lane represents 1 kb Plus DNA Ladder (NEB, Ipswich, MA). DNA integrity is maintained in the enzymatically treated DNA.

**Supplemental Figure 4**

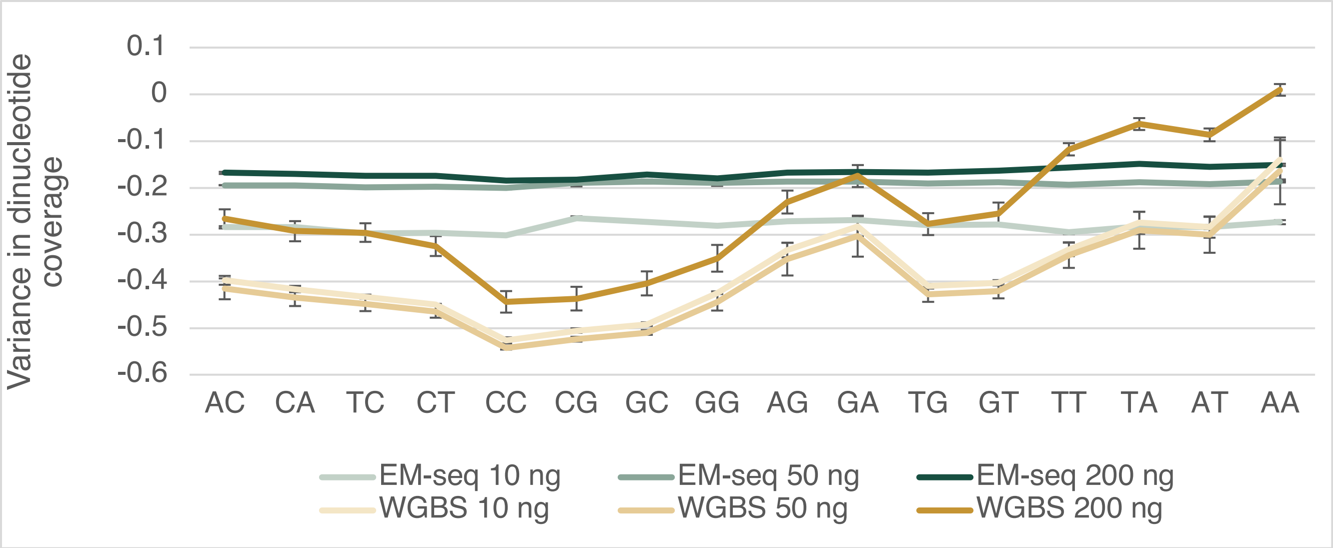


**Nucleotide analysis of EM-seq and Bisulfite Libraries.**

EM-seq and WGBS libraries were made using 10 ng, 50 ng and 200 ng of NA12878 DNA. Dinucleotide distribution for the data was normalized to coverage observed in an unconverted Ultra II DNA library using the same input DNA as the EM-seq and WGBS libraries. The even distribution of dinucleotides found in EM-seq libraries provides evidence for reduced bias compared to WGBS.

**Supplemental Figure 5**

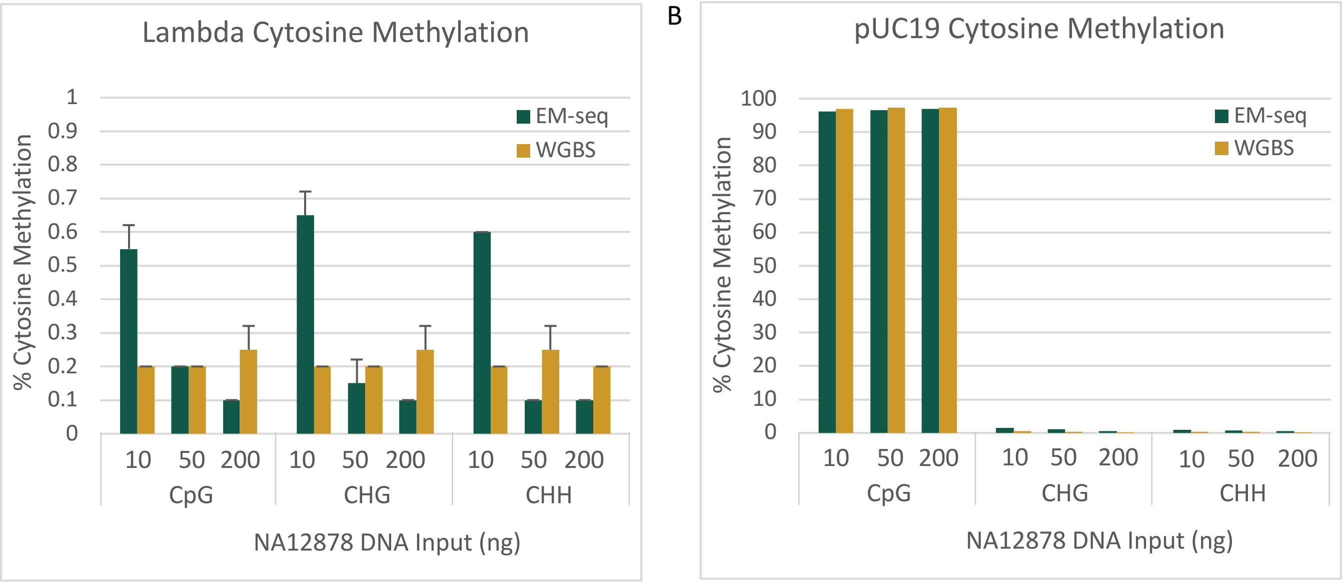


**Cytosine methylation in the EM-seq and bisulfite NA12878 library internal controls**

(A) Methylation of cytosines in the unmethylated lambda controls. EM-seq and WGBS have <1% methylation in CpG, CHG and CHH contexts. (B) Methylation of cytosines in the CpG methylated pUC19 control. EM-seq and WGBS have approximately 97% methylation in the CpG context and around 1% in the CHG and CHH contexts.

#### Supplementary Figure 6

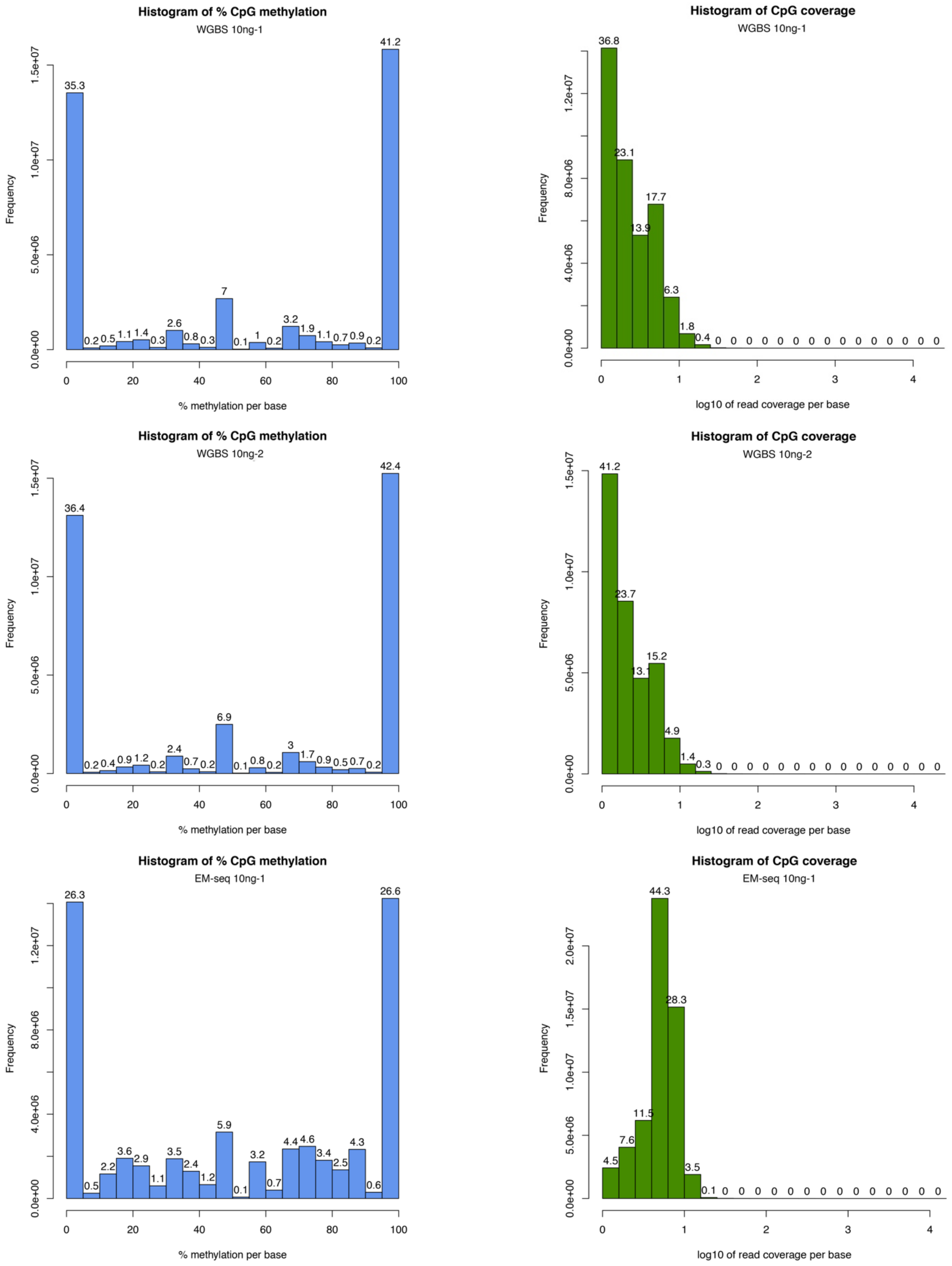

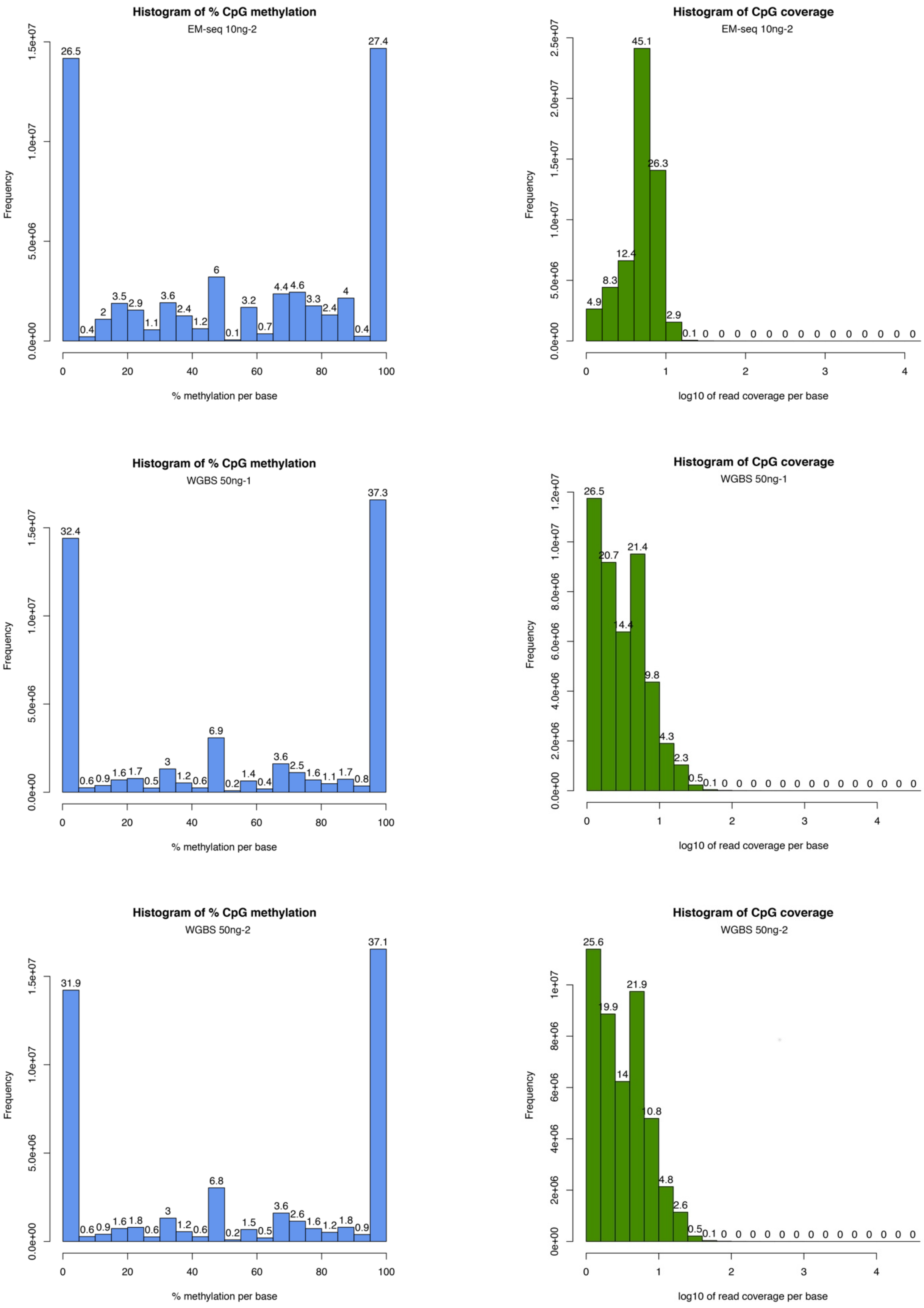
Histogram of % CpG methylation
EM-seq 10ng-2
26.5 0.4 2 3.5 2.9 1.1 3.6 2.4 1.2 6 0.1 3.2 0.7 4.4 4.6 3.3 2.4 4 0.4 27.4
Frequency
% methylation per base
Histogram of CpG coverage
EM-seq 10ng-2
4.9 8.3 12.4 45.1 26.3 2.9 0.1 0 0 0 0 0 0 0 0 0 0 0 0 0 0
Frequency
log10 of read coverage per base
Histogram of % CpG methylation
WGBS 50ng-1
32.4 0.6 0.9 1.6 1.7 0.5 3 1.2 0.6 6.9 0.2 1.4 0.4 3.6 2.5 1.6 1.1 1.7 0.8 37.3
Frequency
% methylation per base
Histogram of CpG coverage
WGBS 50ng-1
26.5 20.7 14.4 21.4 9.8 4.3 2.3 0.5 0.1 0 0 0 0 0 0 0 0 0 0 0 0 0 0
Frequency
log10 of read coverage per base
Histogram of % CpG methylation
WGBS 50ng-2
31.9 0.6 0.9 1.6 1.8 0.6 3 1.2 0.6 6.8 0.2 1.5 0.5 3.6 2.6 1.6 1.2 1.8 0.9 37.1
Frequency
% methylation per base
Histogram of CpG coverage
WGBS 50ng-2
25.6 19.9 14 21.9 10.8 4.8 2.6 0.5 0.1 0 0 0 0 0 0 0 0 0 0 0 0 0 0
Frequency
log10 of read coverage per base

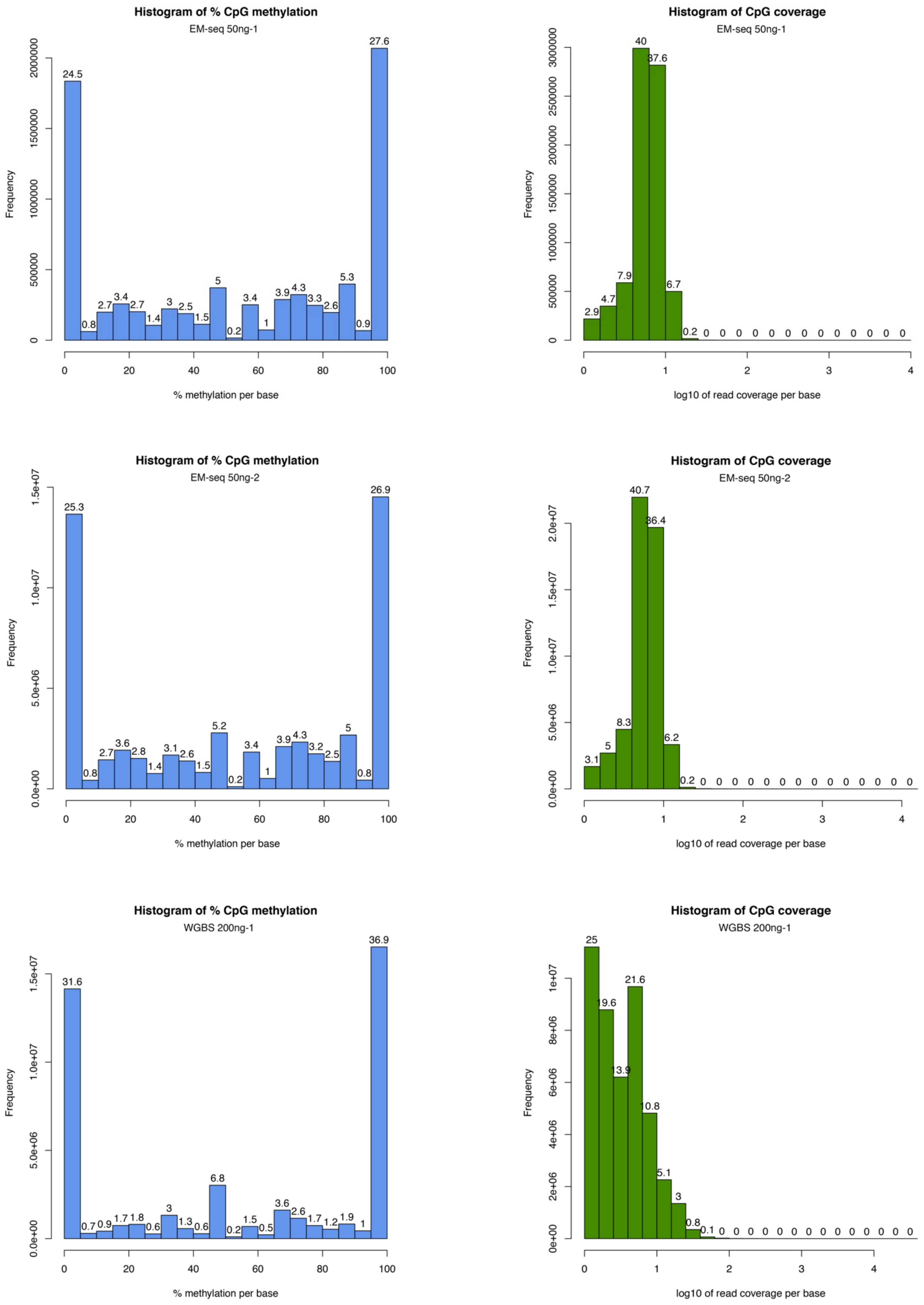

Histogram of % CpG methylation
EM-seq 50ng-1
Frequency
0
500000
1000000
1500000
2000000
24.5 0.8 2.7 3.4 2.7 1.4 3 2.5 1.5 5 0.2 3.4 1 3.9 4.3 3.3 2.6 5.3 0.9 27.6
0 20 40 60 80 100
% methylation per base
Histogram of CpG coverage
EM-seq 50ng-1
Frequency
0
500000
1000000
1500000
2000000
2500000
3000000
2.9 4.7 7.9 40 37.6 6.7 0.2 0 0 0 0 0 0 0 0 0 0 0 0 0 0
0 1 2 3 4
log10 of read coverage per base
Histogram of % CpG methylation
EM-seq 50ng-2
Frequency
0.0e+00
5.0e+06
1.0e+07
1.5e+07
25.3 0.8 2.7 3.6 2.8 1.4 3.1 2.6 1.5 5.2 0.2 3.4 1 3.9 4.3 3.2 2.5 5 0.8 26.9
0 20 40 60 80 100
% methylation per base
Histogram of CpG coverage
EM-seq 50ng-2
Frequency
0.0e+00
5.0e+06
1.0e+07
1.5e+07
2.0e+07
3.1 5 8.3 40.7 36.4 6.2 0.2 0 0 0 0 0 0 0 0 0 0 0 0 0 0
0 1 2 3 4
log10 of read coverage per base
Histogram of % CpG methylation
WGBS 200ng-1
Frequency
0.0e+00
5.0e+06
1.0e+07
1.5e+07
31.6 0.7 0.9 1.7 1.8 0.6 3 1.3 0.6 6.8 0.2 1.5 0.5 3.6 2.6 1.7 1.2 1.9 1 36.9
0 20 40 60 80 100
% methylation per base
Histogram of CpG coverage
WGBS 200ng-1
Frequency
0e+00
2e+06
4e+06
6e+06
8e+06
1e+07
25 19.6 13.9 21.6 10.8 5.1 3 0.8 0.1 0 0 0 0 0 0 0 0 0 0 0 0 0
0 1 2 3 4
log10 of read coverage per base

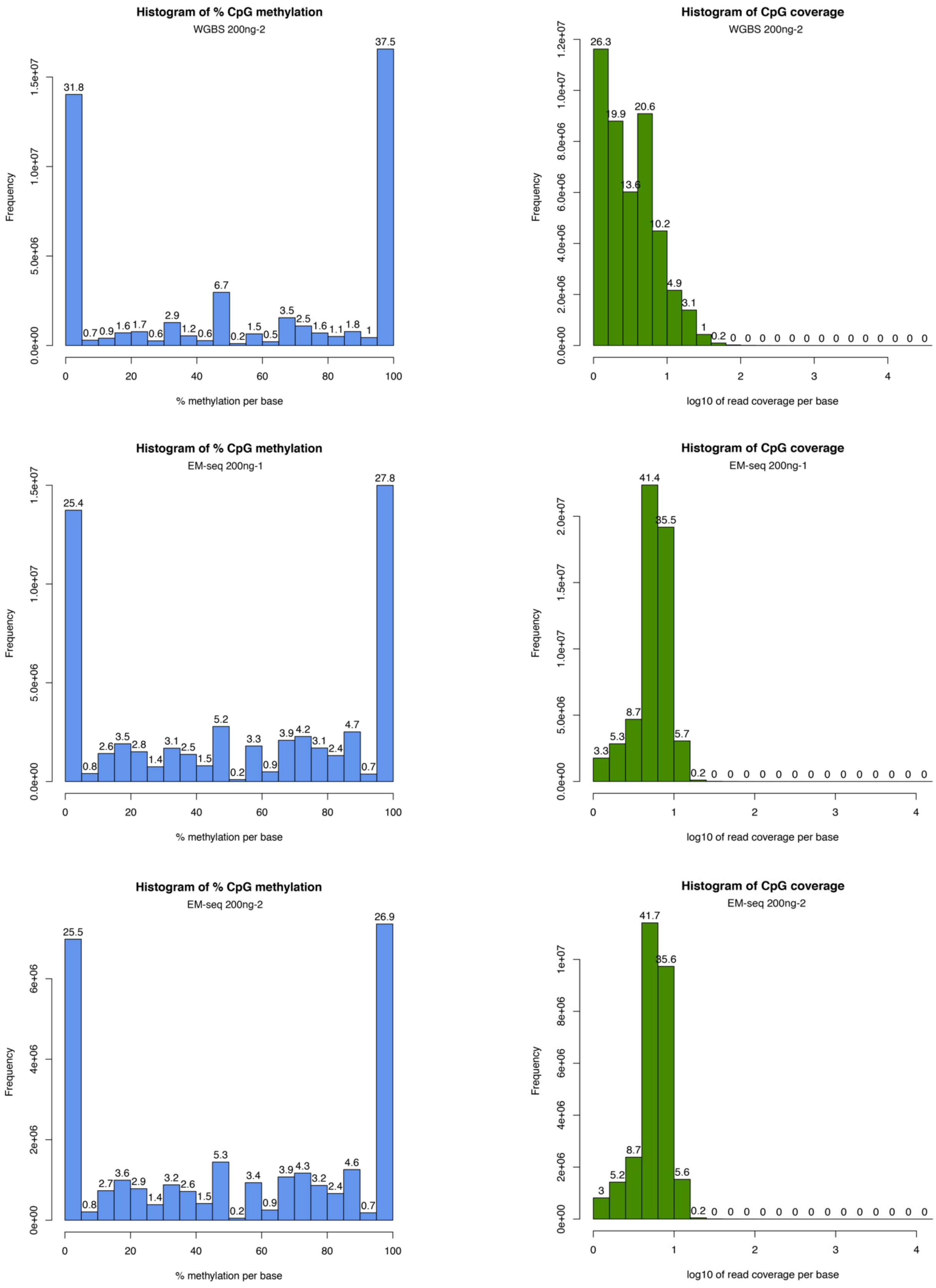

Histogram of % CpG methylation
WGBS 200ng-2
Frequency
% methylation per base
Histogram of CpG coverage
WGBS 200ng-2
Frequency
log10 of read coverage per base
Histogram of % CpG methylation
EM-seq 200ng-1
Frequency
% methylation per base
Histogram of CpG coverage
EM-seq 200ng-1
Frequency
log10 of read coverage per base
Histogram of % CpG methylation
EM-seq 200ng-2
Frequency
% methylation per base
Histogram of CpG coverage
EM-seq 200ng-2
Frequency
log10 of read coverage per base

**CpG Coverage and Methylation Per Base**

Data was analyzed from 10, 50 and 200ng NA12878 EM-seq and bisulfite libraries. CpG coverage and percent methylation per base data were obtained using methylKit.

**Supplemental Figure 7**

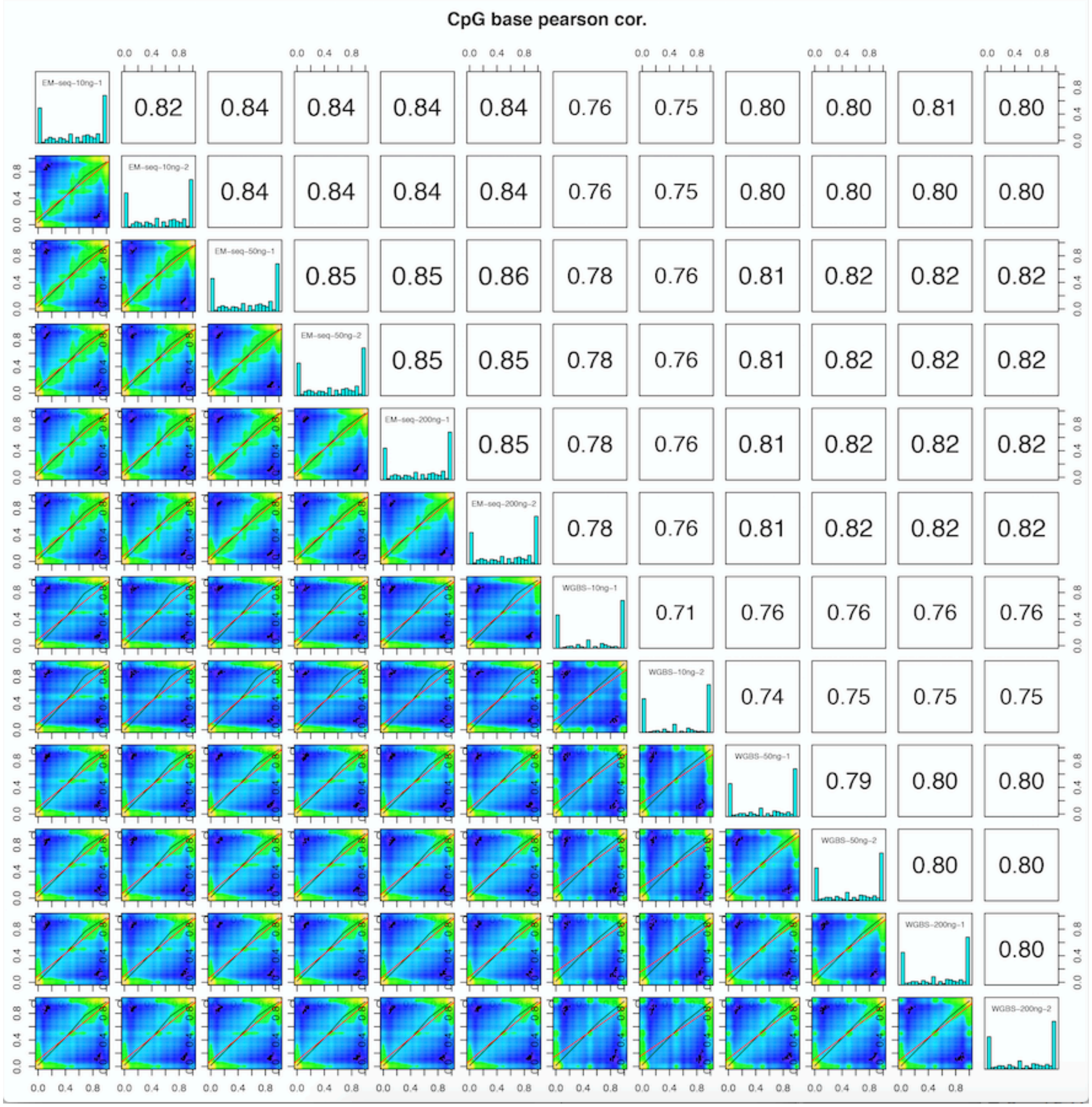


**Correlations between EM-seq and Bisulfite Libraries**

Correlations were plotted using methylKit for 10 ng, 50 ng and 200 ng NA12878 EM-seq and bisulfite libraries at 1x minimum coverage (21 million CpGs common to all libraries). Correlations are highest for EM-seq libraries regardless of input. WGBS library correlations are lower with 10ng inputs driving the lowest correlations and limiting the number of CpGs common to all libraries.

**Supplemental Figure 8**

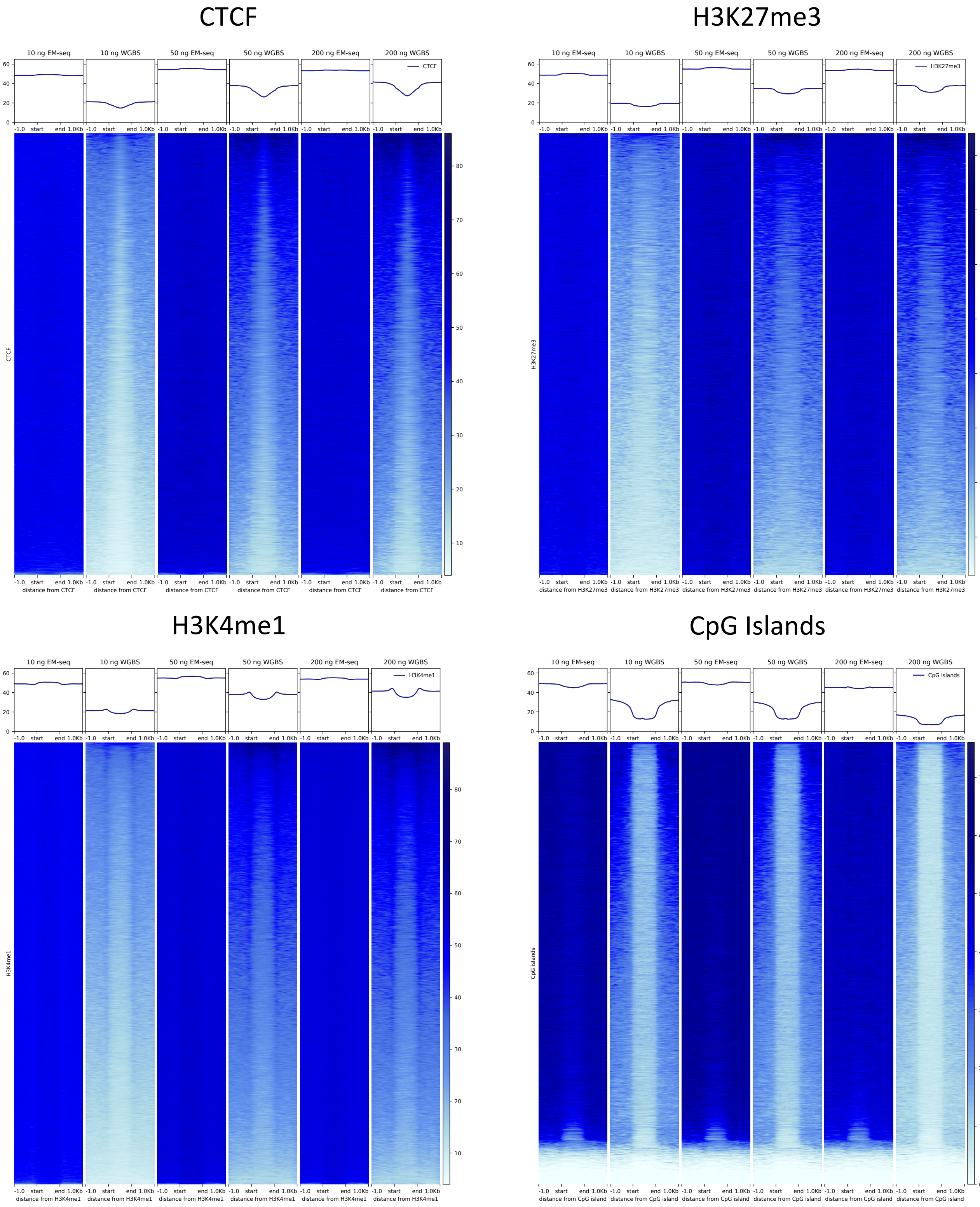

### EZH2

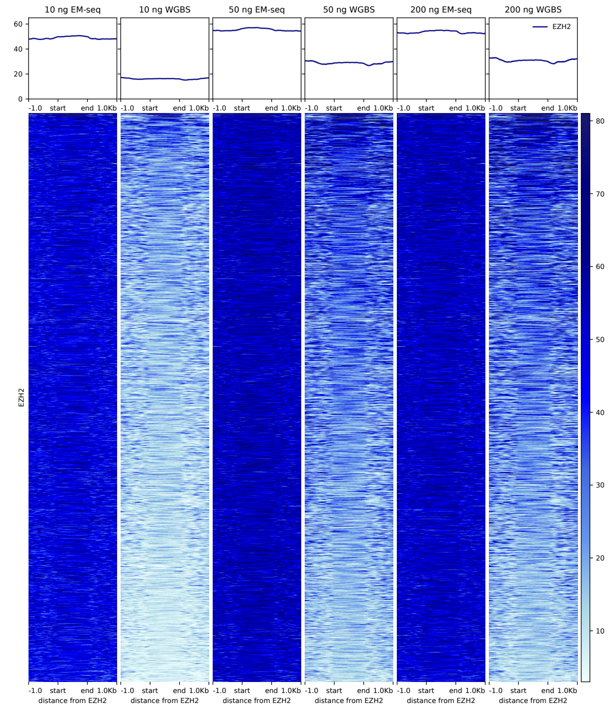


# H3K4me3

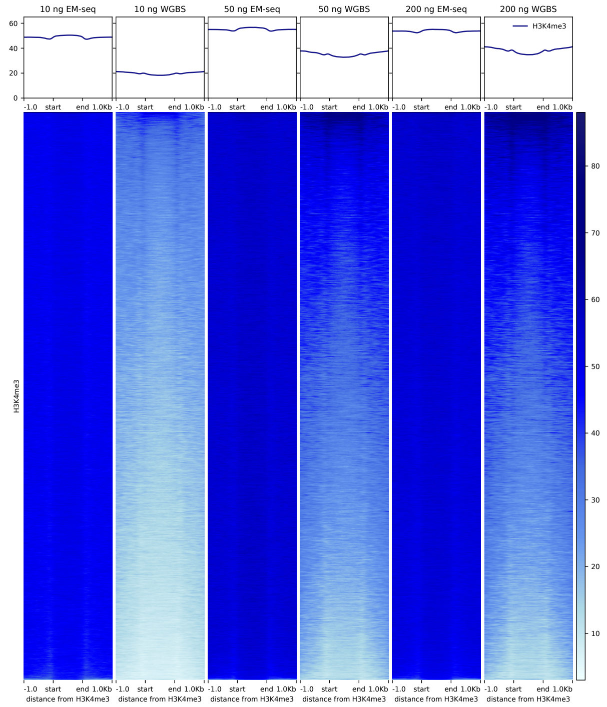


# H3K9ac

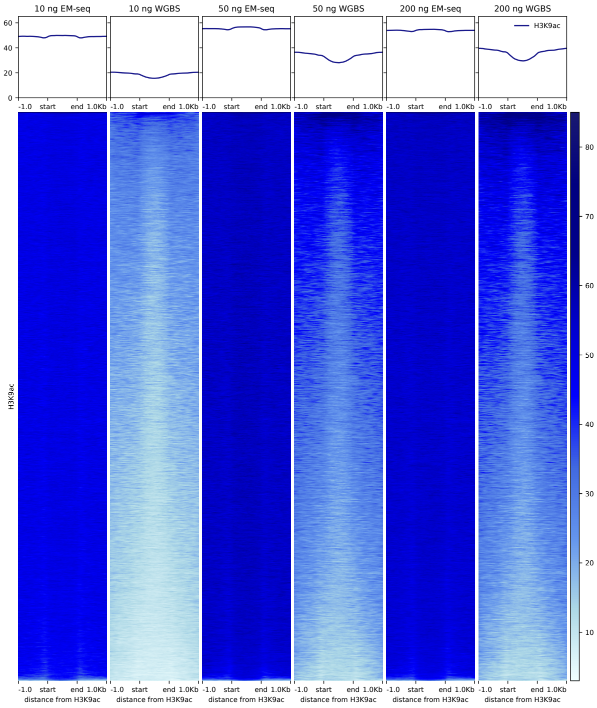


# H3K4me2

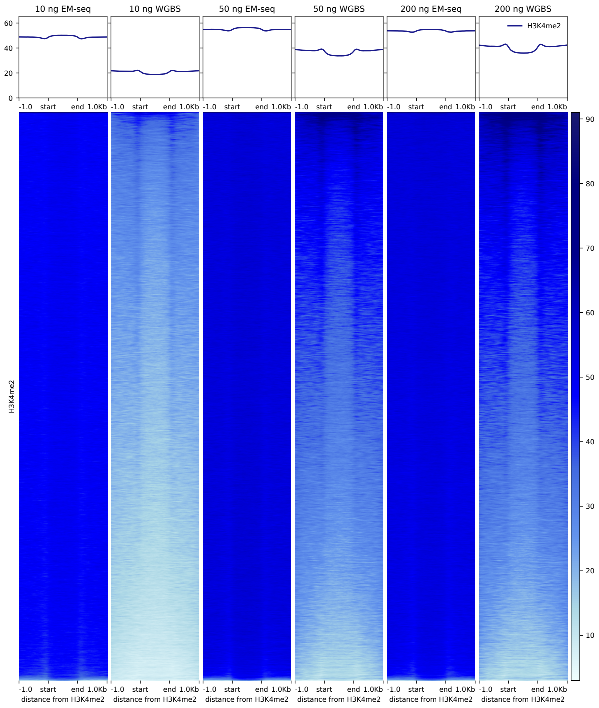

**EM-seq provides more even coverage of CpGs found in diverse genomic features**

Heat maps (made using DeepTools, gencode) representing CpG coverage across a range of genomic features including CTCF and EZH2 binding sites, CpG islands as well as H3K4me1, H3K4me2, H3K4me3, H3K27me3 and H3K9ac binding sites. Coverage plot include +/- 1 kb either side of the start and end of each feature.

**Supplemental Figure 9**

Read Summary (low input)

| Input (ng) | Protocol | Reads | Expected Coverage | % Mapped | % Dups | % Usable |
|---|---|---|---|---|---|---|
| 0.1 | EM-seq (low) | 811M | 26 | 99.8% | 84.5% | 10.8% |
| 0.5 | EM-seq (low) | 811M | 26 | 99.9% | 58.7% | 34.4% |
| 1 | EM-seq (low) | 811M | 26 | 99.9% | 42.6% | 49.8% |
| 10 | Bisulfite (std) | 809M | 26 | 98.2% | 49.9% | 38.9% |
| | EM-seq (low) | 811M | 26 | 100.0% | 13.7% | 77.0% |
| | EM-seq (std) | 811M | 26 | 99.8% | 13.7% | 75.3% |

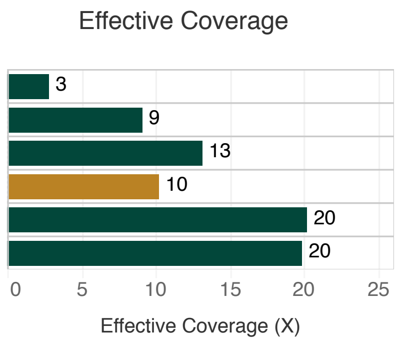


**Sequencing Metrics for Low Input EM-seq Libraries**

EM-seq libraries were made using 100 pg, 500 pg, 1 ng and 10 ng of NA12878 gDNA. Libraries were sequenced on an Illumina NovaSeq 6000, replicates combined, and 810 M total reads per library were used for analysis. Table of sequencing and alignment metrics for EM-seq libraries. At 10 ng inputs, the low input EM-seq method metrics were compared to metrics produced from libraries made using the standard EM-seq method and sodium bisulfite conversion. Metrics were calculated using bwa-meth, Samtools, and Picard. Theoretical coverage is calculated using the number of bases sequenced/total bases in the GRCh38 reference. Percent mapped refers to reads aligned to the reference genome (grch38+controls), Percent Dups refers to reads marked as duplicate by Picard MarkDuplicates, Percent Usable refers to the set of Proper-pair, MapQ 10+, Primary, non-Duplicate reads used in methylation calling (samtools view -F 0xF00 -q 10). Effective Coverage is the Percent Usable * Theoretical coverage.

**Supplemental Figure 10**

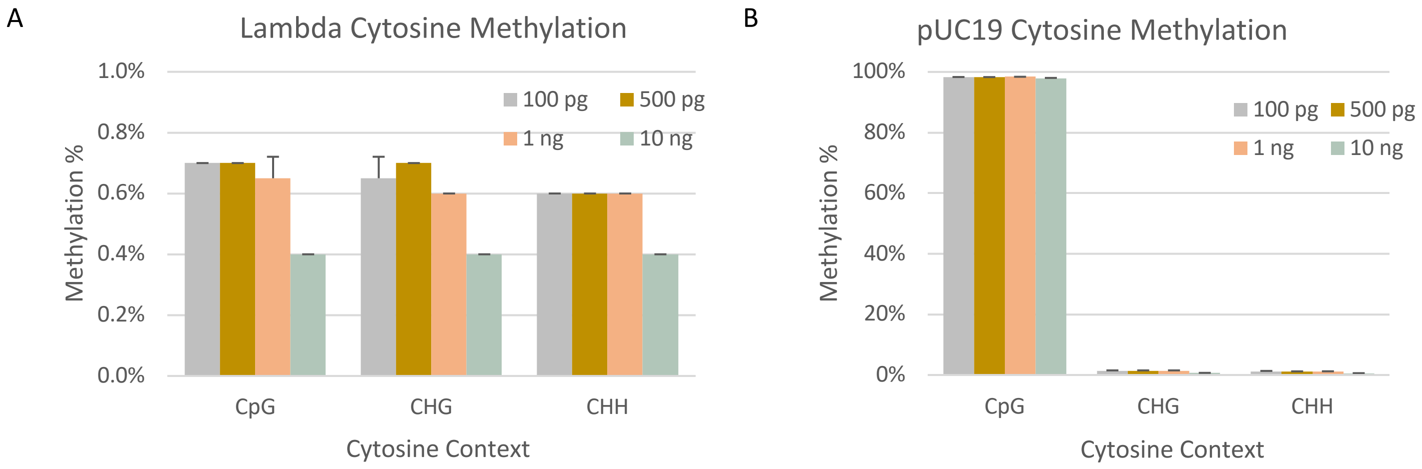


**Low Input EM-seq Libraries, Cytosine Methylation of Lambda and pUC19 internal controls**

(A) Methylation of cytosines in the unmethylated lambda controls. EM-seq has <1% methylation in CpG, CHG and CHH contexts. (B) Methylation of cytosines in the CpG methylated pUC19 control. EM-seq and WGBS have approximately 97% methylation in the CpG context and around 1% in the CHG and CHH contexts.

**Supplemental Figure 11**

A

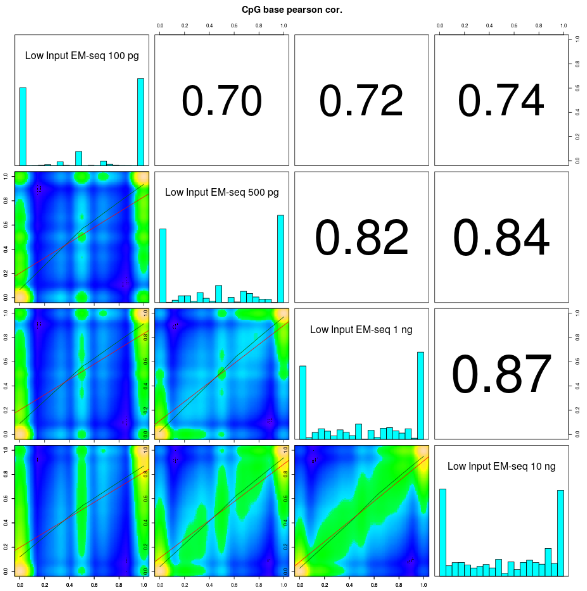


B

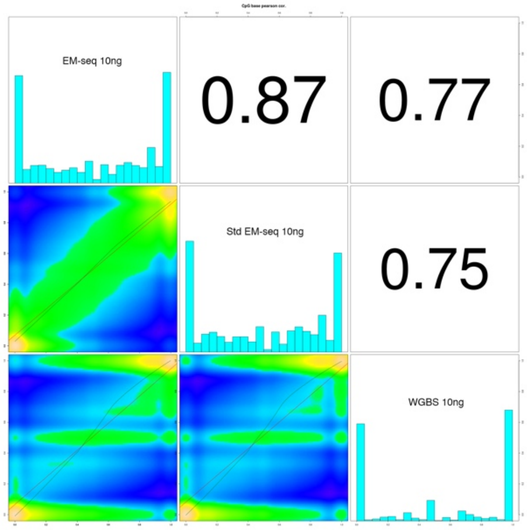


**Correlations between lower DNA input EM-seq Libraries**

Correlations were plotted using methylKit for EM-seq libraries made using NA12878 gDNA. (A) MethylKit correlations for 100 pg, 500 pg, 1 ng, and 10 ng NA12878 EM-seq libraries at 1x minimum coverage. Correlations remain strong between libraries but are reduced when considering the 100 pg DNA input level. (B) Correlations for 10 ng DNA inputs made using methods for low input EM-seq, standard EM-seq and WGBS. Correlation between the two EM-seq methods are higher than when compared to the WGBS method.

**Supplemental Figure 12**

A

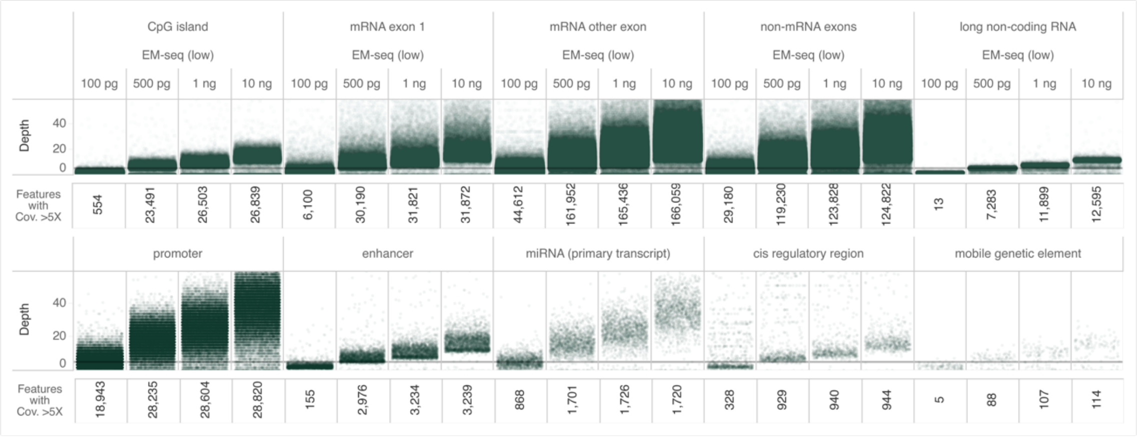


B

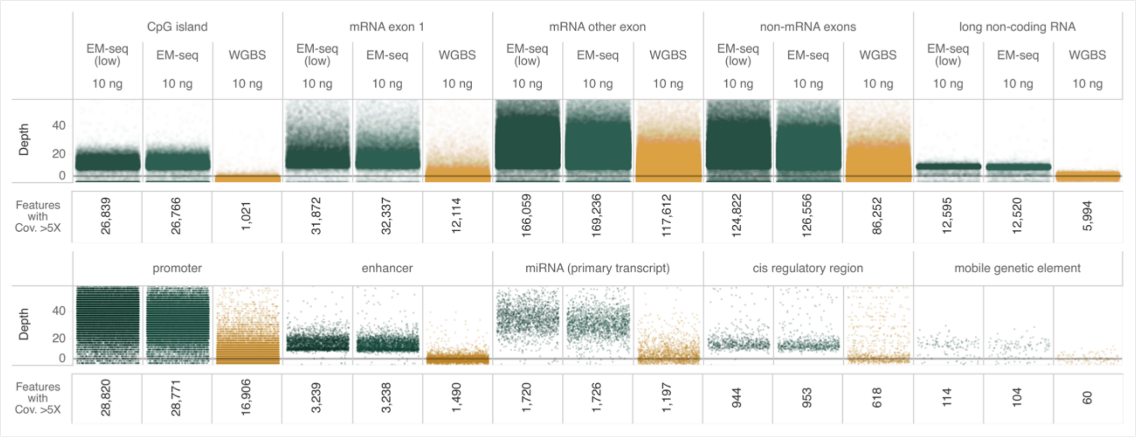


**Low Input EM-seq determines methylation status of cytosines at key genomic features**

Two technical replicates were combined before randomly selecting reads from each library to produce comparable sets of 810 million total human reads. These were analyzed for each library and annotated using featureCounts. (A) The number of genomic features identified at a coverage depth of >5X is similar for 500 pg, 1 ng, and 10 ng DNA inputs. The 100 pg input library has fewer genomic features identified (likely due to lower library complexity at this extreme input amount) (B) Comparison of genomic features identified using 10 ng of NA12878 DNA for low input EM-seq, standard EM-seq and WGBS libraries. A wide range of genomic features at >5X coverage were identified. For EM-seq, the same number of genomic features were identified regardless of low input or standard library preparation method. The number of genomic features in each of the categories was less for WGBS libraries compared to EM-seq libraries.

#### Supplemental Figure 13

A

B

C

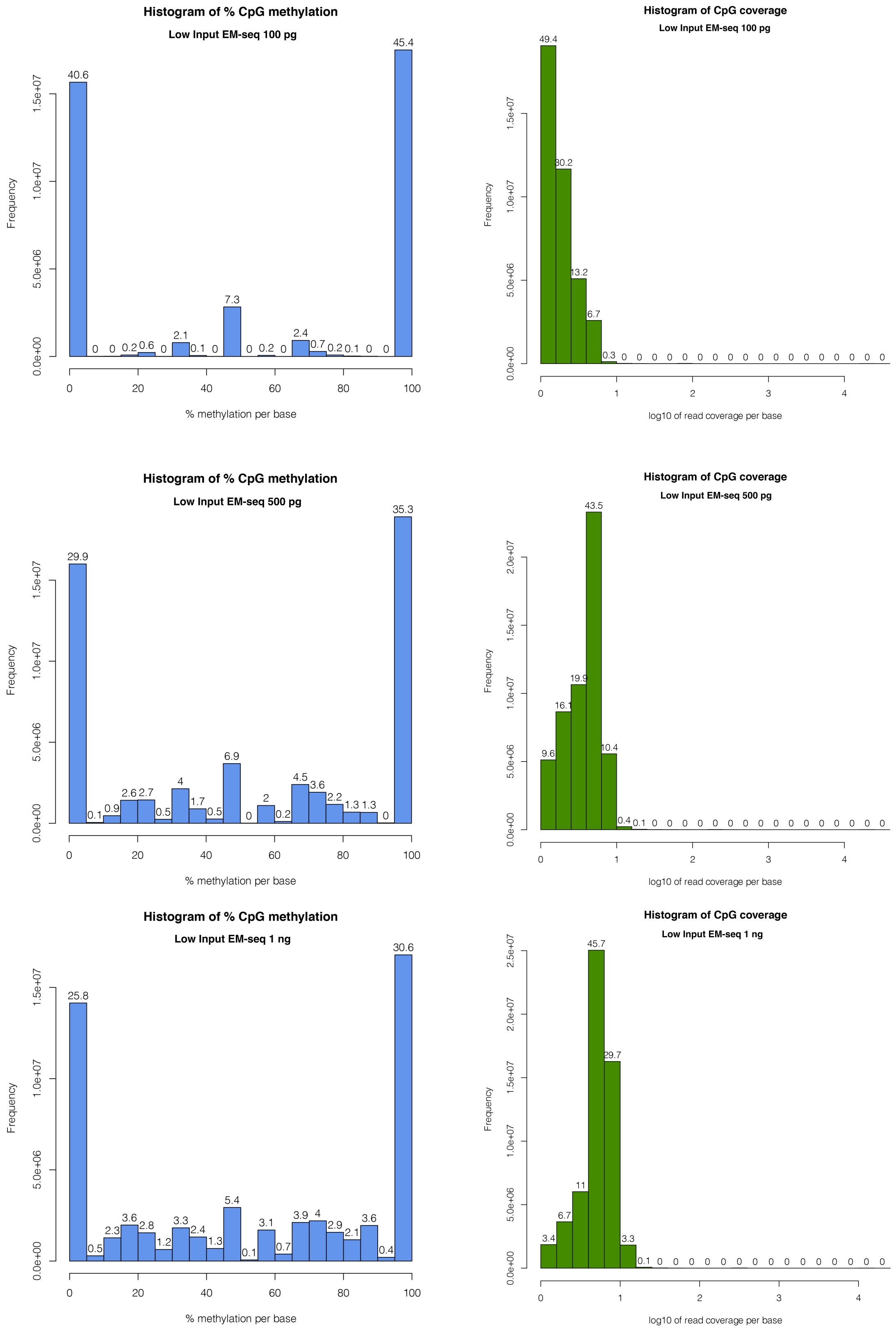

D

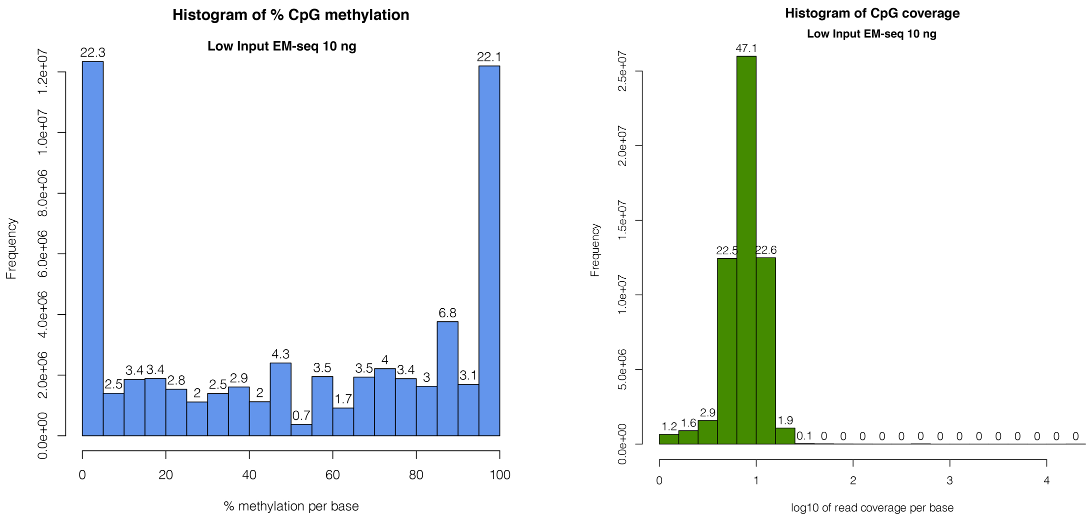


#### CpG coverage and methylation for low input libraries

The coverage and methylation status of CpGs were extracted using MethylDackel and plotting using methylKit for 100 pg, 500 pg, 1 ng and 10 ng NA12878 DNA inputs (A-D). There is deeper coverage for higher DNA inputs. Methylation profiles are as expected, with highest representation of CpG methylation at 0% and 100%.

#### Supplemental Figure 14

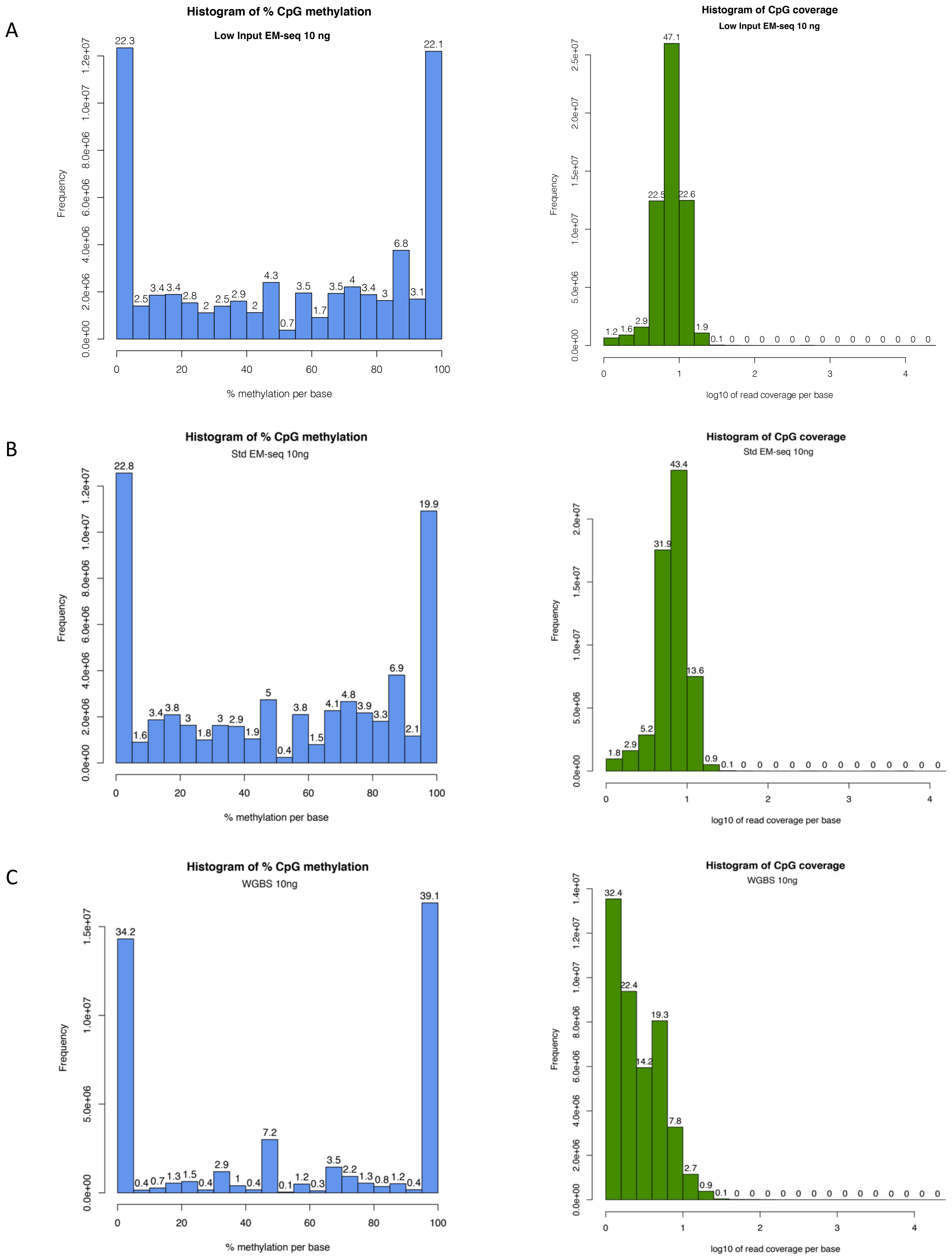

**Comparison of WGBS, standard EM-seq and low input EM-seq 10ng DNA input library types**

The coverage and methylation status of CpGs were extracted from MethylDackel output files using methylKit for 10 ng NA12878 gDNA inputs libraries made using (A) Low input EM-seq methods (B) Standard EM-seq method and (C) WGBS. Methylation profiles and coverage were similar between the two EM-seq methods with highest representation of CpG methylation at 0% and 100%.

**Supplemental Figure 15**

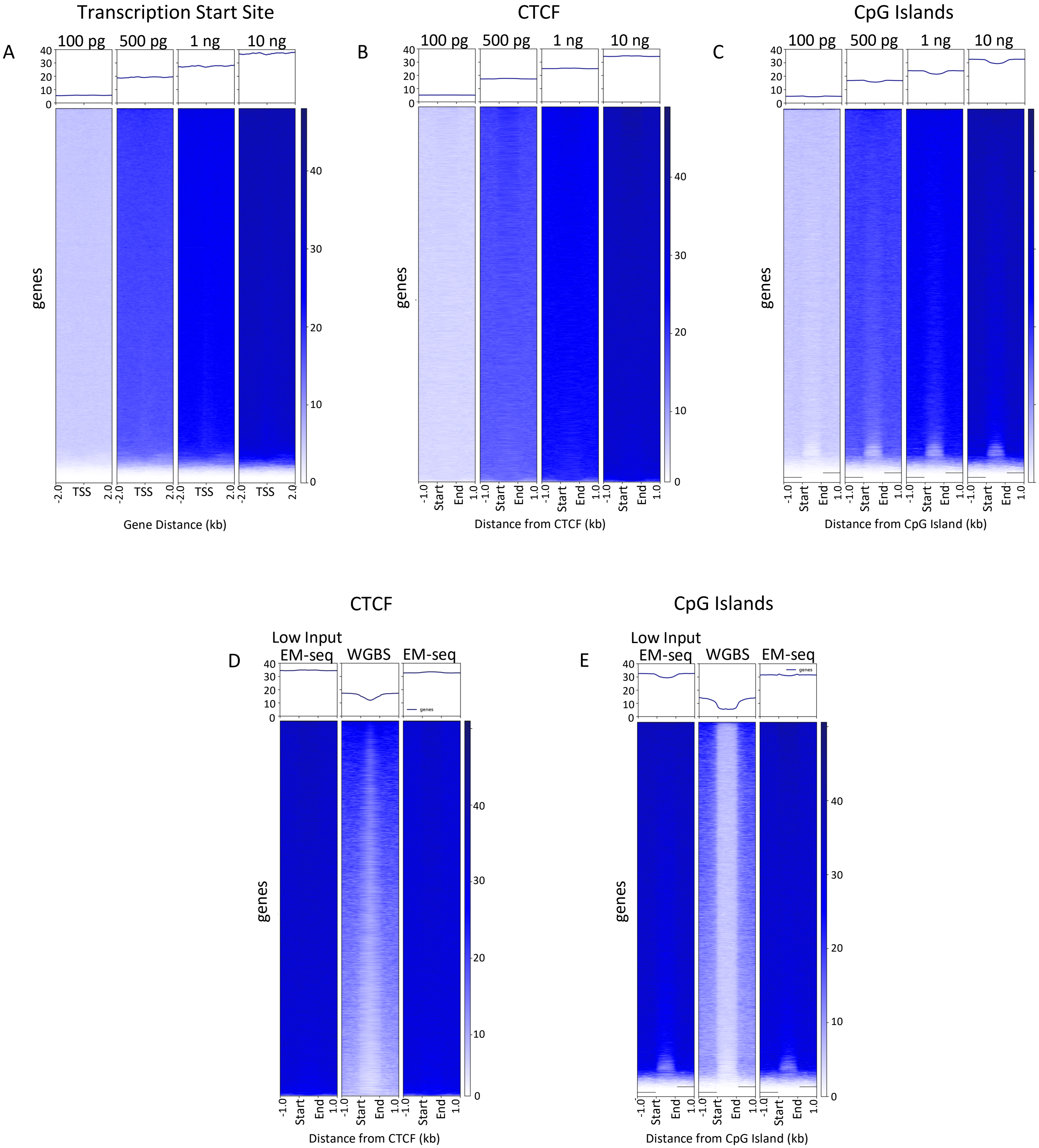

**EM-seq provides greater coverage of CpGs over genomic features**

DeepTools heat maps representing CpG coverage across genomic features including transcription start sites (TSS), CTCF sites, and CpG islands. (A-C) Heat maps for 100 pg, 500 pg, 1 ng and 10 ng DNA inputs showing coverage around TSS, CTCF and CpG islands. Coverage is even across the features at all inputs. (D, E) Heat maps for 10ng DNA inputs comparing coverage of genomic features using libraries prepared using low input EM-seq, standard EM-seq and WGBS. Data from low input and standard EM-seq prepped libraries show similar even coverage. WGBS does not have the same evenness of coverage.
