## Supplemental Tables for "EM-seq: Detection of DNA Methylation at Single Base Resolution from Picograms of DNA"

**Supplemental Table 1**

| Pre-nucleotide substrate | Variable Nucleotide | Oligonucleotide Sequence |
|---|---|---|
| A | C | ATAAGAATAGAATGAAACGTGAAATGAATATGAAATGAATAGTA |
| G | C | ATAAGAATAGAATGAAGCGTGAAATGAATATGAAATGAATAGTA |
| T | C | ATAAGAATAGAATGAATCGTGAAATGAATATGAAATGAATAGTA |
| U | C | ATAAGAATAGAATGAAUCGTGAAATGAATATGAAATGAATAGTA |
| 5mC | C | ATAAGAATAGAATGAA/5mC/CGTGAAATGAATATGAAATGAATAGTA |
| A | 5mC | ATAAGAATAGAATGAAA/5mC/GTGAAATGAATATGAAATGAATAGTA |
| G | 5mC | ATAAGAATAGAATGAAG/5mC/GTGAAATGAATATGAAATGAATAGTA |
| T | 5mC | ATAAGAATAGAATGAAT/5mC/GTGAAATGAATATGAAATGAATAGTA |
| U | 5mC | ATAAGAATAGAATGAAU/5mC/GTGAAATGAATATGAAATGAATAGTA |
| C | 5mC | ATAAGAATAGAATGAAC/5mC/GTGAAATGAATATGAAATGAATAGTA |

**Quantitative analysis of APOBEC3A: site preference specificity oligonucleotides**

Oligonucleotides used in Figure 3A and 3B. For quantitative analysis of APOBEC3A site preference, the same core 44 bp oligonucleotide sequence for investigating the APOBEC3A substrate specificity was used. The base preceding cytosine and 5mC is underlined in the table.

**Supplemental Table 2**

| Variable Nucleotide | Oligonucleotide sequence |
| --- | --- |
| C | ATAAGAATAGAATGAATCGTGAAATGAATATGAAATGAATAGTA |
| 5mC | ATAAGAATAGAATGAAT/5mC/GTGAAATGAATATGAAATGAATAGTA |
| 5hmC | ATAAGAATAGAATGAAT/5hmC/GTGAAATGAATATGAAATGAATAGTA |
| 5gmC | ATAAGAATAGAATGAAT/5gmC/GTGAAATGAATATGAAATGAATAGTA |
| 5fC | ATAAGAATAGAATGAAT/5fC/GTGAAATGAATATGAAATGAATAGTA |
| 5caC | ATAAGAATAGAATGAAT/5caC/GTGAAATGAATATGAAATGAATAGTA |

**Quantitative analysis of APOBEC3A: substrate specificity oligonucleotides**

Oligonucleotides used in Figure 3C. For quantitative analysis of APOBEC3A substrate specificity, 44 bp oligonucleotides were used. These contained the same core sequence, but varied at one single position with either C, 5mC, 5hmC, 5gmC, 5fC or 5caC.

**Supplemental Table 3**

| **Forward primer (5' to 3')** | **Chromosome location (Fwd)** | **Reverse primer (5' to 3')** | **Chromosome location (Rev)** | **Amplicon size, bp** |
|---|---|---|---|---|
| GTAAAATGTTTTGGGTAAAAGTTATATGAATG | Chr1-36100629 | TAAAATAATCCTAACTTCTAACCTCTCTTTTC | Chr1-36097684 | 2945 |
| TGATGAGCGGTAGTTAGGTGAAAG | Chr12-110944577 | AATAAATACAATCCCAAAACTATCAAATCTTC | Chr12-110942370 | 2207 |
| AAAACGGAAATAGGGGTAATGATAG | Chr12-110944120 | AATAAATACAATCCCAAAACTATCAAATCTTC | Chr12-110942370 | 1750 |
| TTTTTTATGATTTGTTGTGATTTGGTTATTAAG | Chr12-110943551 | AATAAATACAATCCCAAAACTATCAAATCTTC | Chr12-110942370 | 1181 |
| GAGTATGGTAAAGTTGAAAATATGTAGATAAG | Chr12-110942913 | AATAAATACAATCCCAAAACTATCAAATCTTC | Chr12-110942370 | 543 |

**DNA integrity assessment**
Sequences of primers used for amplicon test designed using GRCh37 as the reference genome.

**Supplemental Table 4**

| Sample Description | From [bp] | To [bp] | Average Size [bp] | Region Molarity [nmol/l] | PCR cycles |
|---|---|---|---|---|---|
| 0.1ng-1 | 100 | 1000 | 397 | 18.5 | 14 |
| 0.1ng-2 | 100 | 1000 | 382 | 11.3 | 14 |
| 0.5ng-1 | 100 | 1000 | 389 | 13 | 12 |
| 0.5ng-2 | 100 | 1000 | 393 | 14.1 | 12 |
| 1ng-1 | 100 | 1000 | 397 | 12.5 | 11 |
| 1ng-2 | 100 | 1000 | 385 | 11.9 | 11 |
| 10ng-1 | 100 | 1000 | 384 | 21.8 | 8 |
| 10ng-2 | 100 | 1000 | 404 | 24.2 | 8 |

NA12878 low input libraries. The average size of the libraries, PCR yields and PCR cycles are indicated.

**Supplemental Table 5**

### Reference genome sequences

| |
|---|
| >chr1 AC:CM000663.2 gi:568336023 LN:248956422 rl:Chromosome M5:6aef897c3d6ff0c78aff06ac189178dd AS:GRCh38 |
| >chr2 AC:CM000664.2 gi:568336022 LN:242193529 rl:Chromosome M5:f98db672eb0993dcfdabafe2a882905c AS:GRCh38 |
| >chr3 AC:CM000665.2 gi:568336021 LN:198295559 rl:Chromosome M5:76635a41ea913a405ded820447d067b0 AS:GRCh38 |
| >chr4 AC:CM000666.2 gi:568336020 LN:190214555 rl:Chromosome M5:3210fecf1eb92d5489da4346b3fddc6e AS:GRCh38 |
| >chr5 AC:CM000667.2 gi:568336019 LN:181538259 rl:Chromosome M5:a811b3dc9fe66af729dc0dddf7fa4f13 AS:GRCh38 hm:47309185-49591369 |
| >chr6 AC:CM000668.2 gi:568336018 LN:170805979 rl:Chromosome M5:5691468a67c7e7a7b5f2a3a683792c29 AS:GRCh38 |
| >chr7 AC:CM000669.2 gi:568336017 LN:159345973 rl:Chromosome M5:cc044cc2256a1141212660fb07b6171e AS:GRCh38 |
| >chr8 AC:CM000670.2 gi:568336016 LN:145138636 rl:Chromosome M5:c67955b5f7815a9a1edfaa15893d3616 AS:GRCh38 |
| >chr9 AC:CM000671.2 gi:568336015 LN:138394717 rl:Chromosome M5:6c198acf68b5af7b9d676dfdd531b5de AS:GRCh38 |
| >chr10 AC:CM000672.2 gi:568336014 LN:133797422 rl:Chromosome M5:c0eeee7acfdaf31b770a509bdaa6e51a AS:GRCh38 |
| >chr11 AC:CM000673.2 gi:568336013 LN:135086622 rl:Chromosome M5:1511375dc2dd1b633af8cf439ae90cec AS:GRCh38 |
| >chr12 AC:CM000674.2 gi:568336012 LN:133275309 rl:Chromosome M5:96e414eace405d8c27a6d35ba19df56f AS:GRCh38 |
| >chr13 AC:CM000675.2 gi:568336011 LN:114364328 rl:Chromosome M5:a5437debe2ef9c9ef8f3ea2874ae1d82 AS:GRCh38 |
| >chr14 AC:CM000676.2 gi:568336010 LN:107043718 rl:Chromosome M5:e0f0eecc3bcab6178c62b6211565c807 AS:GRCh38 hm:multiple |
| >chr15 AC:CM000677.2 gi:568336009 LN:101991189 rl:Chromosome M5:f036bd11158407596ca6bf3581454706 AS:GRCh38 |
| >chr16 AC:CM000678.2 gi:568336008 LN:90338345 rl:Chromosome M5:db2d37c8b7d019caaf2dd64ba3a6f33a AS:GRCh38 |
| >chr17 AC:CM000679.2 gi:568336007 LN:83257441 rl:Chromosome M5:f9a0fb01553adb183568e3eb9d8626db AS:GRCh38 |
| >chr18 AC:CM000680.2 gi:568336006 LN:80373285 rl:Chromosome M5:11eeaa801f6b0e2e36a1138616b8ee9a AS:GRCh38 |
| >chr19 AC:CM000681.2 gi:568336005 LN:58617616 rl:Chromosome M5:85f9f4fc152c58cb7913c06d6b98573a AS:GRCh38 hm:multiple |
| >chr20 AC:CM000682.2 gi:568336004 LN:64444167 rl:Chromosome M5:b18e6c531b0bd70e949a7fc20859cb01 AS:GRCh38 |
| >chr21 AC:CM000683.2 gi:568336003 LN:46709983 rl:Chromosome M5:974dc7aec0b755b19f031418fdedf293 AS:GRCh38 hm:multiple |
| >chr22 AC:CM000684.2 gi:568336002 LN:50818468 rl:Chromosome M5:ac37ec46683600f808cdd41eac1d55cd AS:GRCh38 hm:multiple |
| >chrX AC:CM000685.2 gi:568336001 LN:156040895 rl:Chromosome M5:2b3a55ff7f58eb308420c8a9b11cac50 AS:GRCh38 |
| >chrY AC:CM000686.2 gi:568336000 LN:57227415 rl:Chromosome M5:ce3e31103314a704255f3cd90369ecce AS:GRCh38 hm:10001-2781479,56887903-57217415 |
| >chrM AC:J01415.2 gi:113200490 LN:16569 rl:Mitochondrion M5:c68f52674c9fb33aef52dcf399755519 AS:GRCh38 tp:circular |
| >chr1_KI270706v1_random AC:KI270706.1 gi:568335410 LN:175055 rg:chr1 rl:unlocalized M5:62def1a794b3e18192863d187af956e6 AS:GRCh38 |
| >chr1_KI270707v1_random AC:KI270707.1 gi:568335409 LN:32032 rg:chr1 rl:unlocalized M5:78135804eb15220565483b7cdd02f3be AS:GRCh38 |
| >chr1_KI270708v1_random AC:KI270708.1 gi:568335408 LN:127682 rg:chr1 rl:unlocalized M5:1e95e047b98ed92148dd84d6c037158c AS:GRCh38 |
| >chr1_KI270709v1_random AC:KI270709.1 gi:568335407 LN:66860 rg:chr1 rl:unlocalized M5:4e2db2933ea96aee8dab54af60ecb37d AS:GRCh38 |

| |
|---|
| >chr1_KI270710v1_random AC:KI270710.1 gi:568335406 LN:40176 rg:chr1 rl:unlocalized<br>M5:9949f776680c6214512ee738ac5da289 AS:GRCh38 |
| >chr1_KI270711v1_random AC:KI270711.1 gi:568335405 LN:42210 rg:chr1 rl:unlocalized<br>M5:af383f98cf4492c1f1c4e750c26cbb40 AS:GRCh38 |
| >chr1_KI270712v1_random AC:KI270712.1 gi:568335404 LN:176043 rg:chr1 rl:unlocalized<br>M5:c38a0fecae6a1838a405406f724d6838 AS:GRCh38 |
| >chr1_KI270713v1_random AC:KI270713.1 gi:568335403 LN:40745 rg:chr1 rl:unlocalized<br>M5:cb78d48cc0adbc58822a1c6fe89e3569 AS:GRCh38 |
| >chr1_KI270714v1_random AC:KI270714.1 gi:568335402 LN:41717 rg:chr1 rl:unlocalized<br>M5:42f7a452b8b769d051ad738ee9f00631 AS:GRCh38 |
| >chr2_KI270715v1_random AC:KI270715.1 gi:568335401 LN:161471 rg:chr2 rl:unlocalized<br>M5:b65a8af1d7bbb7f3c77eea85423452bb AS:GRCh38 |
| >chr2_KI270716v1_random AC:KI270716.1 gi:568335400 LN:153799 rg:chr2 rl:unlocalized<br>M5:2828e63b8edc5e845bf48e75fbad2926 AS:GRCh38 |
| >chr3_GL000221v1_random AC:GL000221.1 gi:224183270 LN:155397 rg:chr3 rl:unlocalized<br>M5:3238fb74ea87ae857f9c7508d315babb AS:GRCh38 |
| >chr4_GL000008v2_random AC:GL000008.2 gi:568335399 LN:209709 rg:chr4 rl:unlocalized<br>M5:a999388c587908f80406444cebe80ba3 AS:GRCh38 |
| >chr5_GL000208v1_random AC:GL000208.1 gi:224183050 LN:92689 rg:chr5 rl:unlocalized<br>M5:aa81be49bf3fe63a79bdc6a6f279abf6 AS:GRCh38 |
| >chr9_KI270717v1_random AC:KI270717.1 gi:568335398 LN:40062 rg:chr9 rl:unlocalized<br>M5:796773a1ee67c988b4de887addbed9e7 AS:GRCh38 |
| >chr9_KI270718v1_random AC:KI270718.1 gi:568335397 LN:38054 rg:chr9 rl:unlocalized<br>M5:b0c463c8efa8d64442b48e936368dad5 AS:GRCh38 |
| >chr9_KI270719v1_random AC:KI270719.1 gi:568335396 LN:176845 rg:chr9 rl:unlocalized<br>M5:cd5e932cfc4c74d05bb64e2126873a3a AS:GRCh38 |
| >chr9_KI270720v1_random AC:KI270720.1 gi:568335395 LN:39050 rg:chr9 rl:unlocalized<br>M5:8c2683400a4aeeb40abff96652b9b127 AS:GRCh38 |
| >chr11_KI270721v1_random AC:KI270721.1 gi:568335394 LN:100316 rg:chr11 rl:unlocalized<br>M5:9654b5d3f36845bb9d19a6dbd15d2f22 AS:GRCh38 |
| >chr14_GL000009v2_random AC:GL000009.2 gi:568335393 LN:201709 rg:chr14 rl:unlocalized<br>M5:862f555045546733591ff7ab15bcecbe AS:GRCh38 |
| >chr14_GL000225v1_random AC:GL000225.1 gi:224183274 LN:211173 rg:chr14 rl:unlocalized<br>M5:63945c3e6962f28ffd469719a747e73c AS:GRCh38 |
| >chr14_KI270722v1_random AC:KI270722.1 gi:568335392 LN:194050 rg:chr14 rl:unlocalized<br>M5:51f46c9093929e6edc3b4dfd50d803fc AS:GRCh38 |
| >chr14_GL000194v1_random AC:GL000194.1 gi:224183213 LN:191469 rg:chr14 rl:unlocalized<br>M5:6ac8f815bf8e845bb3031b73f812c012 AS:GRCh38 |
| >chr14_KI270723v1_random AC:KI270723.1 gi:568335391 LN:38115 rg:chr14 rl:unlocalized<br>M5:74a4b480675592095fb0c577c515b5df AS:GRCh38 |
| >chr14_KI270724v1_random AC:KI270724.1 gi:568335390 LN:39555 rg:chr14 rl:unlocalized<br>M5:c3fcb15dddf45f91ef7d94e2623ce13b AS:GRCh38 |

| |
|---|
| >chr14_KI270725v1_random AC:KI270725.1 gi:568335389 LN:172810 rg:chr14 rl:unlocalized M5:edc6402e58396b90b8738a5e37bf773d AS:GRCh38 |
| >chr14_KI270726v1_random AC:KI270726.1 gi:568335388 LN:43739 rg:chr14 rl:unlocalized M5:fbe54a3197e2b469ccb2f4b161cfbe86 AS:GRCh38 |
| >chr15_KI270727v1_random AC:KI270727.1 gi:568335387 LN:448248 rg:chr15 rl:unlocalized M5:84fe18a7bf03f3b7fc76cbac8eb583f1 AS:GRCh38 |
| >chr16_KI270728v1_random AC:KI270728.1 gi:568335386 LN:1872759 rg:chr16 rl:unlocalized M5:369ff74cf36683b3066a2ca929d9c40d AS:GRCh38 |
| >chr17_GL000205v2_random AC:GL000205.2 gi:568335385 LN:185591 rg:chr17 rl:unlocalized M5:458e71cd53dd1df4083dc7983a6c82c4 AS:GRCh38 |
| >chr17_KI270729v1_random AC:KI270729.1 gi:568335384 LN:280839 rg:chr17 rl:unlocalized M5:2756f6ee4f5780acce31e995443508b6 AS:GRCh38 |
| >chr17_KI270730v1_random AC:KI270730.1 gi:568335383 LN:112551 rg:chr17 rl:unlocalized M5:48f98ede8e28a06d241ab2e946c15e07 AS:GRCh38 |
| >chr22_KI270731v1_random AC:KI270731.1 gi:568335382 LN:150754 rg:chr22 rl:unlocalized M5:8176d9a20401e8d9f01b7ca8b51d9c08 AS:GRCh38 |
| >chr22_KI270732v1_random AC:KI270732.1 gi:568335381 LN:41543 rg:chr22 rl:unlocalized M5:d837bab5e416450df6e1038ae6cd0817 AS:GRCh38 |
| >chr22_KI270733v1_random AC:KI270733.1 gi:568335380 LN:179772 rg:chr22 rl:unlocalized M5:f1fa05d48bb0c1f87237a28b66f0be0b AS:GRCh38 |
| >chr22_KI270734v1_random AC:KI270734.1 gi:568335379 LN:165050 rg:chr22 rl:unlocalized M5:1d17410ae2569c758e6dd51616412d32 AS:GRCh38 |
| >chr22_KI270735v1_random AC:KI270735.1 gi:568335378 LN:42811 rg:chr22 rl:unlocalized M5:eb6b07b73dd9a47252098ed3d9fb78b8 AS:GRCh38 |
| >chr22_KI270736v1_random AC:KI270736.1 gi:568335377 LN:181920 rg:chr22 rl:unlocalized M5:2ff189f33cfa52f321accddf648c5616 AS:GRCh38 |
| >chr22_KI270737v1_random AC:KI270737.1 gi:568335376 LN:103838 rg:chr22 rl:unlocalized M5:2ea8bc113a8193d1d700b584b2c5f42a AS:GRCh38 |
| >chr22_KI270738v1_random AC:KI270738.1 gi:568335375 LN:99375 rg:chr22 rl:unlocalized M5:854ec525c7b6a79e7268f515b6a9877c AS:GRCh38 |
| >chr22_KI270739v1_random AC:KI270739.1 gi:568335374 LN:73985 rg:chr22 rl:unlocalized M5:760fbd73515fedcc9f37737c4a722d6a AS:GRCh38 |
| >chrY_KI270740v1_random AC:KI270740.1 gi:568335373 LN:37240 rg:chrY rl:unlocalized M5:69e42252aead509bf56f1ea6fda91405 AS:GRCh38 |
| >chrUn_KI270302v1 AC:KI270302.1 gi:568335372 LN:2274 rl:unplaced M5:ee6dff38036f7d03478c70717643196e AS:GRCh38 |
| >chrUn_KI270304v1 AC:KI270304.1 gi:568335371 LN:2165 rl:unplaced M5:9423c1b46a48aa6331a77ab5c702ac9d AS:GRCh38 |
| >chrUn_KI270303v1 AC:KI270303.1 gi:568335370 LN:1942 rl:unplaced M5:2cb746c78e0faa11e628603a4bc9bd58 AS:GRCh38 |
| >chrUn_KI270305v1 AC:KI270305.1 gi:568335369 LN:1472 rl:unplaced M5:f9acea3395b6992cf3418b6689b98cf9 AS:GRCh38 |
| >chrUn_KI270322v1 AC:KI270322.1 gi:568335368 LN:21476 rl:unplaced M5:7d459255d1c54369e3b64e719061a5a5 AS:GRCh38 |
| >chrUn_KI270320v1 AC:KI270320.1 gi:568335367 LN:4416 rl:unplaced M5:d898b9c5a0118e76481bf5695272959e AS:GRCh38 |
| >chrUn_KI270310v1 AC:KI270310.1 gi:568335366 LN:1201 rl:unplaced M5:af6cb123af7007793bac06485a2a20e9 AS:GRCh38 |
| >chrUn_KI270316v1 AC:KI270316.1 gi:568335365 LN:1444 rl:unplaced M5:6adde7a9fe7bd6918f12d0f0924aa8ba AS:GRCh38 |

| |
|---|
| >chrUn_KI270315v1 AC:KI270315.1 gi:568335364 LN:2276 rl:unplaced M5:ecc43e822dc011fae1fcfd9981c46e9c AS:GRCh38 |
| >chrUn_KI270312v1 AC:KI270312.1 gi:568335363 LN:998 rl:unplaced M5:26499f2fe4c65621fd8f4ecafbad31d7 AS:GRCh38 |
| >chrUn_KI270311v1 AC:KI270311.1 gi:568335362 LN:12399 rl:unplaced M5:59594f9012d8ce21ed5d1119c051a2ba AS:GRCh38 |
| >chrUn_KI270317v1 AC:KI270317.1 gi:568335361 LN:37690 rl:unplaced M5:cd4b1fda800f6ec9ea8001994dbf6499 AS:GRCh38 |
| >chrUn_KI270412v1 AC:KI270412.1 gi:568335360 LN:1179 rl:unplaced M5:7bb9612f733fb7f098be79499d46350c AS:GRCh38 |
| >chrUn_KI270411v1 AC:KI270411.1 gi:568335359 LN:2646 rl:unplaced M5:fc240322d91d43c04f349cc58fda3eca AS:GRCh38 |
| >chrUn_KI270414v1 AC:KI270414.1 gi:568335358 LN:2489 rl:unplaced M5:753e02ef3b1c590e0e3376ad2ebb5836 AS:GRCh38 |
| >chrUn_KI270419v1 AC:KI270419.1 gi:568335357 LN:1029 rl:unplaced M5:58455e7a788f0dc82034d1fb109f6f5c AS:GRCh38 |
| >chrUn_KI270418v1 AC:KI270418.1 gi:568335356 LN:2145 rl:unplaced M5:1537ec12b9c58d137a2d4cb9db896bbc AS:GRCh38 |
| >chrUn_KI270420v1 AC:KI270420.1 gi:568335355 LN:2321 rl:unplaced M5:bac954a897539c91982a7e3985a49910 AS:GRCh38 |
| >chrUn_KI270424v1 AC:KI270424.1 gi:568335354 LN:2140 rl:unplaced M5:747c8f94f34d5de4ad289bc604708210 AS:GRCh38 |
| >chrUn_KI270417v1 AC:KI270417.1 gi:568335353 LN:2043 rl:unplaced M5:cd26758fda713f9c96e51d541f50c2d0 AS:GRCh38 |
| >chrUn_KI270422v1 AC:KI270422.1 gi:568335352 LN:1445 rl:unplaced M5:3fce80eb4c0554376b591699031feb56 AS:GRCh38 |
| >chrUn_KI270423v1 AC:KI270423.1 gi:568335351 LN:981 rl:unplaced M5:bdf5a85c001731dccfb150db2bfe58ac AS:GRCh38 |
| >chrUn_KI270425v1 AC:KI270425.1 gi:568335350 LN:1884 rl:unplaced M5:665a46879bbb48294b0cfa87b61e71f6 AS:GRCh38 |
| >chrUn_KI270429v1 AC:KI270429.1 gi:568335349 LN:1361 rl:unplaced M5:ee8962dbef9396884f649e566b78bf06 AS:GRCh38 |
| >chrUn_KI270442v1 AC:KI270442.1 gi:568335348 LN:392061 rl:unplaced M5:796289c4cda40e358991f9e672490015 AS:GRCh38 |
| >chrUn_KI270466v1 AC:KI270466.1 gi:568335347 LN:1233 rl:unplaced M5:530b7033716a5d72dd544213c513fd12 AS:GRCh38 |
| >chrUn_KI270465v1 AC:KI270465.1 gi:568335346 LN:1774 rl:unplaced M5:bb1b2323414425c46531b3c3d22ae00d AS:GRCh38 |
| >chrUn_KI270467v1 AC:KI270467.1 gi:568335345 LN:3920 rl:unplaced M5:db34e0dc109a4afd499b5ec6aaae9754 AS:GRCh38 |
| >chrUn_KI270435v1 AC:KI270435.1 gi:568335344 LN:92983 rl:unplaced M5:1655c75415b9c29e143a815f44286d05 AS:GRCh38 |
| >chrUn_KI270438v1 AC:KI270438.1 gi:568335343 LN:112505 rl:unplaced M5:e765271939b854dd6826aa764e930c87 AS:GRCh38 |
| >chrUn_KI270468v1 AC:KI270468.1 gi:568335342 LN:4055 rl:unplaced M5:0a603090f06108ed7aff75df0767b822 AS:GRCh38 |
| >chrUn_KI270510v1 AC:KI270510.1 gi:568335341 LN:2415 rl:unplaced M5:cd7348b3b5d9d0dfef6aed2af75ce920 AS:GRCh38 |
| >chrUn_KI270509v1 AC:KI270509.1 gi:568335340 LN:2318 rl:unplaced M5:1cdeb8c823d839e1d1735b5bc9a14856 AS:GRCh38 |
| >chrUn_KI270518v1 AC:KI270518.1 gi:568335339 LN:2186 rl:unplaced M5:3fd898b62ca859f50fb8b83e7706352b AS:GRCh38 |
| >chrUn_KI270508v1 AC:KI270508.1 gi:568335338 LN:1951 rl:unplaced M5:7d42a358d472b9cbdfdf30c8742473d0 AS:GRCh38 |
| >chrUn_KI270516v1 AC:KI270516.1 gi:568335337 LN:1300 rl:unplaced M5:1cbaaafbbf016906a5bf5886f5a0ecb7 AS:GRCh38 |
| >chrUn_KI270512v1 AC:KI270512.1 gi:568335336 LN:22689 rl:unplaced M5:ba1021c82d1230af856f59079e2f71b4 AS:GRCh38 |
| >chrUn_KI270519v1 AC:KI270519.1 gi:568335335 LN:138126 rl:unplaced M5:8d754e9c9afd904fba0a2cd577fcc9a1 AS:GRCh38 |
| >chrUn_KI270522v1 AC:KI270522.1 gi:568335334 LN:5674 rl:unplaced M5:070b4678e22501029c2e3297115216bc AS:GRCh38 |
| >chrUn_KI270511v1 AC:KI270511.1 gi:568335333 LN:8127 rl:unplaced M5:907ca34a4a2a6673632ebaf513a4c1a4 AS:GRCh38 |
| >chrUn_KI270515v1 AC:KI270515.1 gi:568335332 LN:6361 rl:unplaced M5:dd7527ee8e0bdb0a43ca0b2a5456c8c3 AS:GRCh38 |
| >chrUn_KI270507v1 AC:KI270507.1 gi:568335331 LN:5353 rl:unplaced M5:311894d0a815eb07c5cac49da851cb4a AS:GRCh38 |
| >chrUn_KI270517v1 AC:KI270517.1 gi:568335330 LN:3253 rl:unplaced M5:913440c38d95c618617ca69bb9296170 AS:GRCh38 |

| |
|---|
| >chrUn_KI270529v1 AC:KI270529.1 gi:568335329 LN:1899 rl:unplaced M5:4caf890f2586daab8e4b2e2db904f05f AS:GRCh38 |
| >chrUn_KI270528v1 AC:KI270528.1 gi:568335328 LN:2983 rl:unplaced M5:d75c9235f0b8c449fc4352997c56b086 AS:GRCh38 |
| >chrUn_KI270530v1 AC:KI270530.1 gi:568335327 LN:2168 rl:unplaced M5:04549369e1197c626669a10164613635 AS:GRCh38 |
| >chrUn_KI270539v1 AC:KI270539.1 gi:568335326 LN:993 rl:unplaced M5:19e3a982e67eafef39c5a3e4163f1e17 AS:GRCh38 |
| >chrUn_KI270538v1 AC:KI270538.1 gi:568335325 LN:91309 rl:unplaced M5:d60b72221cc7af871f2c757577e4c92a AS:GRCh38 |
| >chrUn_KI270544v1 AC:KI270544.1 gi:568335324 LN:1202 rl:unplaced M5:e62a14b14467cdf48b5a236c66918f0f AS:GRCh38 |
| >chrUn_KI270548v1 AC:KI270548.1 gi:568335323 LN:1599 rl:unplaced M5:866b0db8e9cf66208c2c064bd09ce0a2 AS:GRCh38 |
| >chrUn_KI270583v1 AC:KI270583.1 gi:568335322 LN:1400 rl:unplaced M5:b127e2e6dbe358ff192b271b8c6ee690 AS:GRCh38 |
| >chrUn_KI270587v1 AC:KI270587.1 gi:568335321 LN:2969 rl:unplaced M5:36be47659719f47b95caa51744aa8f70 AS:GRCh38 |
| >chrUn_KI270580v1 AC:KI270580.1 gi:568335320 LN:1553 rl:unplaced M5:1df30dae0f605811d927dcea58e729fc AS:GRCh38 |
| >chrUn_KI270581v1 AC:KI270581.1 gi:568335319 LN:7046 rl:unplaced M5:9f26945f9f9b3f865c9ebe953cbbc1a9 AS:GRCh38 |
| >chrUn_KI270579v1 AC:KI270579.1 gi:568335318 LN:31033 rl:unplaced M5:fe62fb1964002717cc1b034630e89b1f AS:GRCh38 |
| >chrUn_KI270589v1 AC:KI270589.1 gi:568335317 LN:44474 rl:unplaced M5:211c215414693fe0a2399cf82e707e03 AS:GRCh38 |
| >chrUn_KI270590v1 AC:KI270590.1 gi:568335316 LN:4685 rl:unplaced M5:e8a57f147561b361091791b9010cd28b AS:GRCh38 |
| >chrUn_KI270584v1 AC:KI270584.1 gi:568335315 LN:4513 rl:unplaced M5:d93636c9d54abd013cfc0d4c01334032 AS:GRCh38 |
| >chrUn_KI270582v1 AC:KI270582.1 gi:568335314 LN:6504 rl:unplaced M5:6fd9804a7478d2e28160fe9f017689cb AS:GRCh38 |
| >chrUn_KI270588v1 AC:KI270588.1 gi:568335313 LN:6158 rl:unplaced M5:37ffa850e69b342a8f8979bd3ffc77d4 AS:GRCh38 |
| >chrUn_KI270593v1 AC:KI270593.1 gi:568335312 LN:3041 rl:unplaced M5:f4a5bfa203e9e81acb640b18fb11e78e AS:GRCh38 |
| >chrUn_KI270591v1 AC:KI270591.1 gi:568335311 LN:5796 rl:unplaced M5:d6af509d69835c9ac25a30086e5a4051 AS:GRCh38 |
| >chrUn_KI270330v1 AC:KI270330.1 gi:568335310 LN:1652 rl:unplaced M5:c2c590706a339007b00c59e0b8937e78 AS:GRCh38 |
| >chrUn_KI270329v1 AC:KI270329.1 gi:568335309 LN:1040 rl:unplaced M5:f023f927ae84c5cc48dc4dce11ba90f2 AS:GRCh38 |
| >chrUn_KI270334v1 AC:KI270334.1 gi:568335308 LN:1368 rl:unplaced M5:53afe12d1371f250a3d1de655345d374 AS:GRCh38 |
| >chrUn_KI270333v1 AC:KI270333.1 gi:568335307 LN:2699 rl:unplaced M5:57baf650c47bba9b3a8b7c6d0fb55ad6 AS:GRCh38 |
| >chrUn_KI270335v1 AC:KI270335.1 gi:568335306 LN:1048 rl:unplaced M5:eb27188639503b524d2659a23b8262ea AS:GRCh38 |
| >chrUn_KI270338v1 AC:KI270338.1 gi:568335305 LN:1428 rl:unplaced M5:301ef75a6b2996d745eb3464bd352b57 AS:GRCh38 |
| >chrUn_KI270340v1 AC:KI270340.1 gi:568335304 LN:1428 rl:unplaced M5:56b462bac20d385cdfcde0155fe4c3a1 AS:GRCh38 |
| >chrUn_KI270336v1 AC:KI270336.1 gi:568335303 LN:1026 rl:unplaced M5:69ad2d85d870c8b0269434581e86e30e AS:GRCh38 |
| >chrUn_KI270337v1 AC:KI270337.1 gi:568335302 LN:1121 rl:unplaced M5:16fc8d71a2662a6cfec7bdeec3d810c6 AS:GRCh38 |
| >chrUn_KI270363v1 AC:KI270363.1 gi:568335301 LN:1803 rl:unplaced M5:6edd17a912f391022edbc192d49f2489 AS:GRCh38 |
| >chrUn_KI270364v1 AC:KI270364.1 gi:568335300 LN:2855 rl:unplaced M5:6ff66a8e589ca27d93b5bac0e5b13a87 AS:GRCh38 |
| >chrUn_KI270362v1 AC:KI270362.1 gi:568335299 LN:3530 rl:unplaced M5:bc82401ffd9a5ae711fa0ea34da8d2f0 AS:GRCh38 |
| >chrUn_KI270366v1 AC:KI270366.1 gi:568335298 LN:8320 rl:unplaced M5:44a0b65b7ba6bcff37eca202e7d966ea AS:GRCh38 |
| >chrUn_KI270378v1 AC:KI270378.1 gi:568335297 LN:1048 rl:unplaced M5:fc13bda7dbd914c92fb7e49489d1350f AS:GRCh38 |
| >chrUn_KI270379v1 AC:KI270379.1 gi:568335296 LN:1045 rl:unplaced M5:3218bef25946cd95de585dfc7750f63b AS:GRCh38 |
| >chrUn_KI270389v1 AC:KI270389.1 gi:568335295 LN:1298 rl:unplaced M5:2c9b08c57c27e714d4d5259fd91b6983 AS:GRCh38 |
| >chrUn_KI270390v1 AC:KI270390.1 gi:568335294 LN:2387 rl:unplaced M5:7a64d89ea14990c16d20f4d6e7283e10 AS:GRCh38 |
| >chrUn_KI270387v1 AC:KI270387.1 gi:568335293 LN:1537 rl:unplaced M5:22a12462264340c25e912b8485cdfa91 AS:GRCh38 |
| >chrUn_KI270395v1 AC:KI270395.1 gi:568335292 LN:1143 rl:unplaced M5:7c03ca4756c1620f318fb189214388d8 AS:GRCh38 |
| >chrUn_KI270396v1 AC:KI270396.1 gi:568335291 LN:1880 rl:unplaced M5:9069bed3c2efe7cc87227d619ad5816f AS:GRCh38 |

| |
|---|
| >chrUn_KI270388v1 AC:KI270388.1 gi:568335290 LN:1216 rl:unplaced M5:76f9f3315fa4b831e93c36cd88196480 AS:GRCh38 |
| >chrUn_KI270394v1 AC:KI270394.1 gi:568335289 LN:970 rl:unplaced M5:d5171e863a3d8f832f0559235987b1e5 AS:GRCh38 |
| >chrUn_KI270386v1 AC:KI270386.1 gi:568335288 LN:1788 rl:unplaced M5:b9b1baaa7abf206f6b70cf31654172db AS:GRCh38 |
| >chrUn_KI270391v1 AC:KI270391.1 gi:568335287 LN:1484 rl:unplaced M5:1fa5cf03b3eac0f1b4a64948fd09de53 AS:GRCh38 |
| >chrUn_KI270383v1 AC:KI270383.1 gi:568335286 LN:1750 rl:unplaced M5:694d75683e4a9554bcc1291edbcaee43 AS:GRCh38 |
| >chrUn_KI270393v1 AC:KI270393.1 gi:568335285 LN:1308 rl:unplaced M5:3724e1d70677d6b5c4bcf17fd40da111 AS:GRCh38 |
| >chrUn_KI270384v1 AC:KI270384.1 gi:568335284 LN:1658 rl:unplaced M5:b06e44ea15d0a57618d6ca7d2e6ac5d2 AS:GRCh38 |
| >chrUn_KI270392v1 AC:KI270392.1 gi:568335283 LN:971 rl:unplaced M5:59b3ca8de65fb171683f8a06d3b4bf0d AS:GRCh38 |
| >chrUn_KI270381v1 AC:KI270381.1 gi:568335282 LN:1930 rl:unplaced M5:2a9297cfd3b3807195ab9ad07e775d99 AS:GRCh38 |
| >chrUn_KI270385v1 AC:KI270385.1 gi:568335281 LN:990 rl:unplaced M5:112a8b1df94ef0498a0bfe2d2ea5cc23 AS:GRCh38 |
| >chrUn_KI270382v1 AC:KI270382.1 gi:568335280 LN:4215 rl:unplaced M5:e7085cdcee6ad62f359744e13d3209fc AS:GRCh38 |
| >chrUn_KI270376v1 AC:KI270376.1 gi:568335279 LN:1136 rl:unplaced M5:59e8fc80b78d62325082334b43dffdba AS:GRCh38 |
| >chrUn_KI270374v1 AC:KI270374.1 gi:568335278 LN:2656 rl:unplaced M5:dbc92c9a92e558946e58b4909ec95dd5 AS:GRCh38 |
| >chrUn_KI270372v1 AC:KI270372.1 gi:568335277 LN:1650 rl:unplaced M5:53a9d5e8fd28bce5da5efcfd9114dbf2 AS:GRCh38 |
| >chrUn_KI270373v1 AC:KI270373.1 gi:568335276 LN:1451 rl:unplaced M5:b174fe53be245a840cd6324e39b88ced AS:GRCh38 |
| >chrUn_KI270375v1 AC:KI270375.1 gi:568335275 LN:2378 rl:unplaced M5:d678250c97e9b94aa390fa46e70a6d83 AS:GRCh38 |
| >chrUn_KI270371v1 AC:KI270371.1 gi:568335274 LN:2805 rl:unplaced M5:a0af3d778dfeb7963e8e6d84c0c54fba AS:GRCh38 |
| >chrUn_KI270448v1 AC:KI270448.1 gi:568335273 LN:7992 rl:unplaced M5:0f40827c265cb813b6e723da6c9b926b AS:GRCh38 |
| >chrUn_KI270521v1 AC:KI270521.1 gi:568335272 LN:7642 rl:unplaced M5:af5bef7cefec7bd7efa729ac6c5be088 AS:GRCh38 |
| >chrUn_GL000195v1 AC:GL000195.1 gi:224183229 LN:182896 rl:unplaced M5:5d9ec007868d517e73543b005ba48535 AS:GRCh38 |
| >chrUn_GL000219v1 AC:GL000219.1 gi:224183268 LN:179198 rl:unplaced M5:f977edd13bac459cb2ed4a5457dba1b3 AS:GRCh38 |
| >chrUn_GL000220v1 AC:GL000220.1 gi:224183269 LN:161802 rl:unplaced M5:fc35de963c57bf7648429e6454f1c9db AS:GRCh38 |
| >chrUn_GL000224v1 AC:GL000224.1 gi:224183273 LN:179693 rl:unplaced M5:d5b2fc04f6b41b212a4198a07f450e20 AS:GRCh38 |
| >chrUn_KI270741v1 AC:KI270741.1 gi:568335271 LN:157432 rl:unplaced M5:86eaea8a15a3950e37442eaaa5c9dc92 AS:GRCh38 |
| >chrUn_GL000226v1 AC:GL000226.1 gi:224183275 LN:15008 rl:unplaced M5:1c1b2cd1fccbc0a99b6a447fa24d1504 AS:GRCh38 |
| >chrUn_GL000213v1 AC:GL000213.1 gi:224183287 LN:164239 rl:unplaced M5:9d424fdcc98866650b58f004080a992a AS:GRCh38 |
| >chrUn_KI270743v1 AC:KI270743.1 gi:568335270 LN:210658 rl:unplaced M5:3b62d9d3100f530d509e4efebd98502c AS:GRCh38 |
| >chrUn_KI270744v1 AC:KI270744.1 gi:568335269 LN:168472 rl:unplaced M5:e90aee46b947ff8c32291a6843fde3f9 AS:GRCh38 |
| >chrUn_KI270745v1 AC:KI270745.1 gi:568335268 LN:41891 rl:unplaced M5:1386fe3de6f82956f2124e19353ff9c1 AS:GRCh38 |
| >chrUn_KI270746v1 AC:KI270746.1 gi:568335267 LN:66486 rl:unplaced M5:c470486a0a858e14aa21d7866f83cc17 AS:GRCh38 |

| |
|---|
| >chrUn_KI270747v1 AC:KI270747.1 gi:568335266 LN:198735 rl:unplaced M5:62375d812ece679c9fd2f3d08d4e22a4<br>AS:GRCh38 |
| >chrUn_KI270748v1 AC:KI270748.1 gi:568335265 LN:93321 rl:unplaced M5:4f6c6ab005c852a4352aa33e7cc88ded<br>AS:GRCh38 |
| >chrUn_KI270749v1 AC:KI270749.1 gi:568335264 LN:158759 rl:unplaced M5:c899a7b4e911d371283f3f4058ca08b7<br>AS:GRCh38 |
| >chrUn_KI270750v1 AC:KI270750.1 gi:568335263 LN:148850 rl:unplaced M5:c022ba92f244b7dc54ea90c4eef4d554<br>AS:GRCh38 |
| >chrUn_KI270751v1 AC:KI270751.1 gi:568335262 LN:150742 rl:unplaced M5:1b758bbdee0e9ca882058d916cba9d29<br>AS:GRCh38 |
| >chrUn_KI270752v1 AC:KI270752.1 gi:568335261 LN:27745 rl:unplaced M5:e0880631848337bd58559d9b1519da63<br>AS:GRCh38 |
| >chrUn_KI270753v1 AC:KI270753.1 gi:568335260 LN:62944 rl:unplaced M5:25075fb2a1ecada67c0eb2f1fe0c7ec9<br>AS:GRCh38 |
| >chrUn_KI270754v1 AC:KI270754.1 gi:568335259 LN:40191 rl:unplaced M5:fe9e16233cecbc244f06f3acff3d03b8<br>AS:GRCh38 |
| >chrUn_KI270755v1 AC:KI270755.1 gi:568335258 LN:36723 rl:unplaced M5:4a7da6a658955bd787af8add3ccb5751<br>AS:GRCh38 |
| >chrUn_KI270756v1 AC:KI270756.1 gi:568335257 LN:79590 rl:unplaced M5:2996b120a5a5e15dab6555f0bf92e374<br>AS:GRCh38 |
| >chrUn_KI270757v1 AC:KI270757.1 gi:568335256 LN:71251 rl:unplaced M5:174c73b60b41d8a1ef0fbaa4b3bdf0d3<br>AS:GRCh38 |
| >chrUn_GL000214v1 AC:GL000214.1 gi:224183298 LN:137718 rl:unplaced M5:46c2032c37f2ed899eb41c0473319a69<br>AS:GRCh38 |
| >chrUn_KI270742v1 AC:KI270742.1 gi:568335255 LN:186739 rl:unplaced M5:2f31c013a4a8301deb8ab7ed1ca1cd99<br>AS:GRCh38 |
| >chrUn_GL000216v2 AC:GL000216.2 gi:568335254 LN:176608 rl:unplaced M5:725009a7e3f5b78752b68afa922c090c<br>AS:GRCh38 |
| >chrUn_GL000218v1 AC:GL000218.1 gi:224183305 LN:161147 rl:unplaced M5:1d708b54644c26c7e01c2dad5426d38c<br>AS:GRCh38 |
| >chrEBV AC:AJ507799.2 gi:86261677 LN:171823 rl:decoy M5:6743bd63b3ff2b5b8985d8933c53290a<br>SP:Human_herpesvirus_4 tp:circular |
| >phage_lambda J02459.1_48502 |
| >plasmid_puc19c L09137.2 Cloning vector pUC19c, complete sequence |
| >phage_T4 168903 |
| >phage_Xp12 65076 |
